# Gamete fusion rapidly reconstitutes a bi-partite transcription factor to block re-fertilization

**DOI:** 10.1101/334953

**Authors:** Aleksandar Vještica, Laura Merlini, Pedro N’kosi, Sophie G Martin

## Abstract

The ploidy cycle, integral to sexual reproduction, requires not only meiosis to halve the number of chromosomes, but also mechanisms that ensure zygotes are formed by exactly two partners^1–5^. During sexual reproduction of the fungal model organism *Schizosaccharomyces pombe*, haploid P- and M-cells normally fuse to form a diploid zygote that immediately enters meiosis^6^. Here, we reveal that fast post-fusion reconstitution of a bi-partite transcription factor actively blocks re-fertilization. We first identify mutants that undergo transient cell fusion involving cytosol exchange but not karyogamy, and show this drives distinct cell fates in the two gametes: The P-partner undergoes lethal, haploid meiosis while the M-cell persists in mating. Consistently, we find that the zygotic transcription that drives meiosis is initiated rapidly only from the P-parental genome, even in wild type cells. This asymmetric gene expression depends on a bi-partite complex formed post-fusion between the nuclear P-cell-specific homeobox protein Pi and a cytosolic M-specific peptide Mi^7^,^8^, which is captured by Pi in the P-nucleus. Zygotic transcription is thus poised to initiate in the P-nucleus as fast as Mi reaches it. The asymmetric nuclear accumulation is inherent to the transcription factor design, and is reconstituted by a pair of synthetic interactors, one localized to the nucleus of one gamete and the other in the cytosol of its partner. Strikingly, imposing a delay in zygotic transcription, by postponing Mi expression or deleting its transcriptional target in the P-genome, leads to zygotes fusing with additional gametes, thus forming polyploids and eventually aneuploid progeny. We further show that the signaling cascade to block re-fertilization shares components with, but bifurcates from, meiotic induction^9–11^. Thus, cytoplasmic connection upon gamete fusion leads to rapid reconstitution of a bi-partite transcription factor in one partner to block re-fertilization and induce meiosis, thus ensuring genome maintenance during sexual reproduction.

---

While studying mutants displaying cell fusion defects (ref.^12^), we discovered a striking post-fusion asymmetry between mating partners. Deleting the p21-activated kinase Pak2 impaired cell fusion, resulting in an increased number of unfused partners and extended lifetime of fusion focus components^13^ at the cell-cell contact site (Fig. 1A, S1A). Unexpectedly, *pak2*Δ mating also produced ∼10% of aberrant asci never seen in wildtype, which appeared to be derived from either three parental cells containing more than four spores (Fig. 1A, type IIIa), or a single cell that underwent sporulation (Fig. 1A, type IIIb). Three lines of evidence revealed that these asci result from meiosis and sporulation taking place in haploid cells.

**Figure 1.**
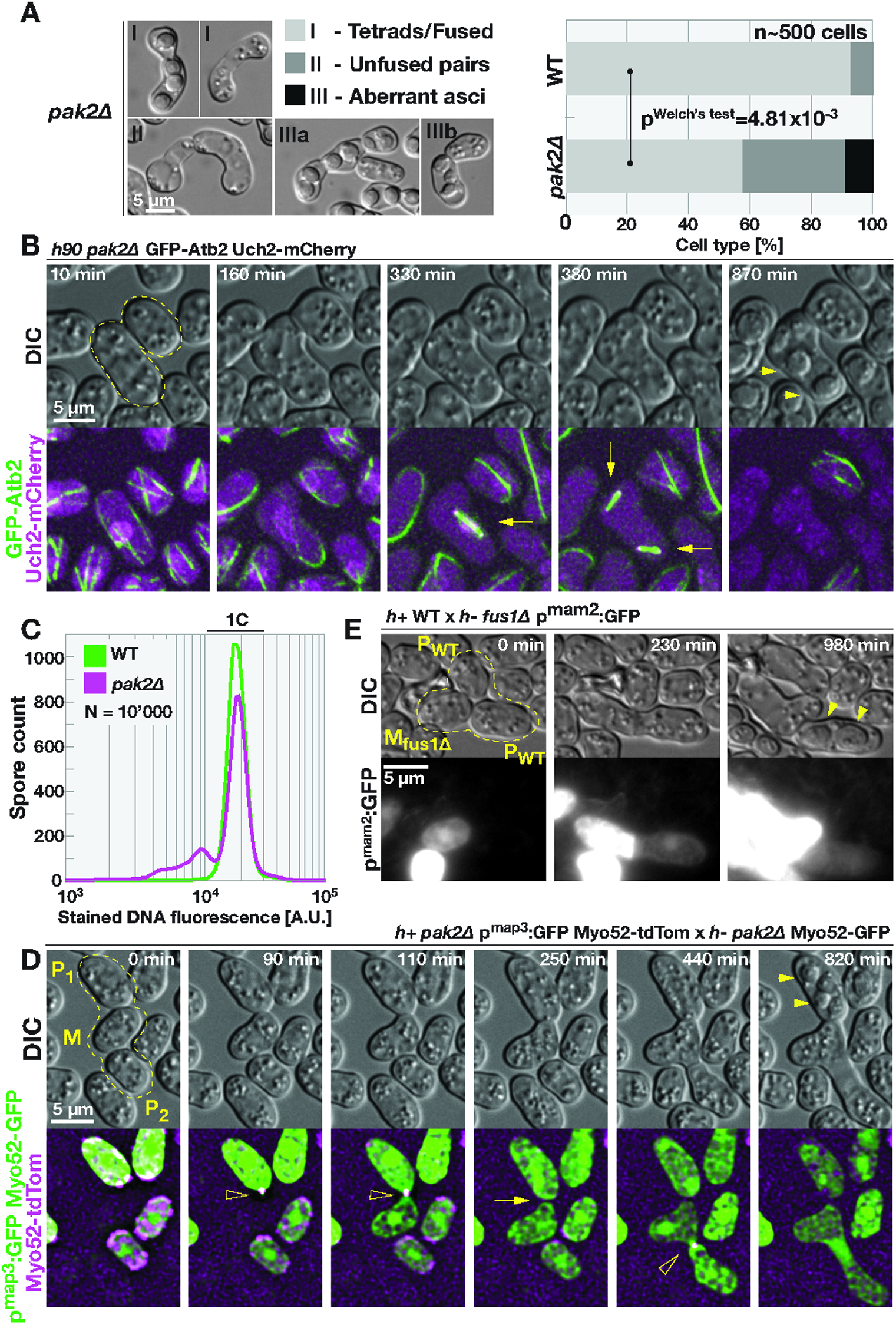
Transient cell fusion induces haploid meiosis and aneuploidy.

First, live imaging showed that karyogamy did not occur between mating partners that produced aberrant asci, even though the spore-forming partner showed microtubule reorganization typical of meiosis: a long cytosolic bundle, characteristic of the meiotic ‘horsetail’ stage^14–17^ followed by two successive rounds of spindle formation, indicative of meiosis-I and meiosis-II (ref.^18^, Fig. 1B, S1B, Mov. S1). The other partner cell maintained interphase microtubule organization and ability to mate with an additional partner (thus yielding type IIIa asci). Second, labeling one chromosomal locus with *lacO*:GFP-NLS-LacI system^19^ revealed that the spore-forming cell contained only two foci distributed between up to four spores, indicating these spores are aneuploid (Fig. S1C). By contrast, wild-type asci formed four spores each with a single focus (ref.^20^, Fig. S1C). Third, flow cytometry analysis of DNA-stained spores from *pak2Δ* crosses indicated presence of spores with less then 1C content (Fig. 1C). We conclude that aberrant *pak2Δ* asci form upon meiosis and sporulation in a single, haploid cell.

Haploid meiosis was provoked by transient fusion between partners: *pak2Δ* cells that produced aberrant asci initially formed a fusion focus and fused, thus exchanging cytosolic GFP, but then re-sealed their fusion pore, as indicated by persistent unequal levels of GFP between partners (n>20, Fig. 1D, S1E, Mov. S2,3). Transient fusion was required to induce haploid meiosis since deleting the Fus1 formin, an essential component of the fusion machinery^13^,^21^, prevented formation of aberrant asci by *pak2Δ* cells (n>200, Fig. S1F). Remarkably, crosses between wild-type and *fus1Δ* mutant, in which fusion efficiency is reduced but not abrogated^13^,^21^, also resulted in transient fusion events each followed by haploid meiosis in one partner, whether this was *fus1*Δ or wildtype (Fig. 1E, S1F-G, Mov. S4). We conclude that transient cell-cell fusion, independently of genotype, induces meiosis in one partner cell.

Quantification of homothallic wildtype and *pak2*Δ mating mixtures for phenotypes described in the main text and represented in inset images. **(B)** Time-lapse of homothallic *pak2*Δ mating cells expressing GFP-α-tubulin in green and nuclear marker Uch2-mCherry in magenta. Note lack of karyogamy in the outlined *pak2*Δ mating pair even though one partner forms meiotic spindles (arrows) and spores (arrowhead), while the other partner maintains interphase microtubule organization. Fig. S1B shows wildtype control. **(C)** Flow cytometry analysis of DNA content in spores produced by wildtype and *pak2*Δ cells. **(D)** Time-lapse of *pak2*Δ mating cells expressing cytosolic GFP from the P-cell specific *p*^*map3*^ promoter. Note the transient cell fusion, observed as exchange of GFP, between M- and P1-cells followed by sealing of the fusion pore (arrow) and spore formation (full arrowheads) in the P1-partner. Also note formation of the fusion focus (empty arrowhead), labeled by Myo52-GFP in the M-cell and Myo52-tdTomato in the P-cell. Fig. S1D reports wildtype control. **(E)** Time-lapse showing transient cell fusion between wildtype P- and *fus1*Δ M-cells expressing GFP from the M-cell specific *p*^*mam2*^ promoter. Persistent GFP signal difference between partners indicated sealing of the fusion pore. Note spore formation (arrowhead) only in the wildtype P-cell.

## Zygotic transcription occurs first in the P-cell

Remarkably, all instances of transient cell fusion induced haploid meiosis strictly in the P-cell (n=64 in *pak2*Δ; n=9 between *fus1*Δ and wild-type), and the master meiotic regulator Mei3^11^,^22^ was induced only in P-cells undergoing transient fusion. In transiently fusing pairs, tagging *mei3* with a fast-folding GFP variant (sfGFP)^23^ in the P-genome resulted in strong fluorescent signal in the P-cell undergoing haploid meiosis, whereas labeling of M-genome *mei3* did not lead to any noticeable fluorescence induction in either partner (Fig. 2A, 2B and Mov. S5, n>10 in each cross). We note that the M-cell encoded Mei3-sfGFP was expressed upon complete fusion that formed normal zygotes (Fig. 2B and Mov. S5, note Mei3-expression in zygote formed by M- and P_2_-cells). Importantly, *mei3* was also asymmetrically expressed in wildtype zygotes: P-genome encoded Mei3-sfGFP expression was significantly more rapid (about 15 min earlier) than Mei3-sfGFP induction from the M-cell genome (Fig. 2C, 2E, S2A and Mov. S6). Thus, fission yeast zygotes express the meiotic inducer Mei3 first from the P-cell genome.

**Figure 2.**
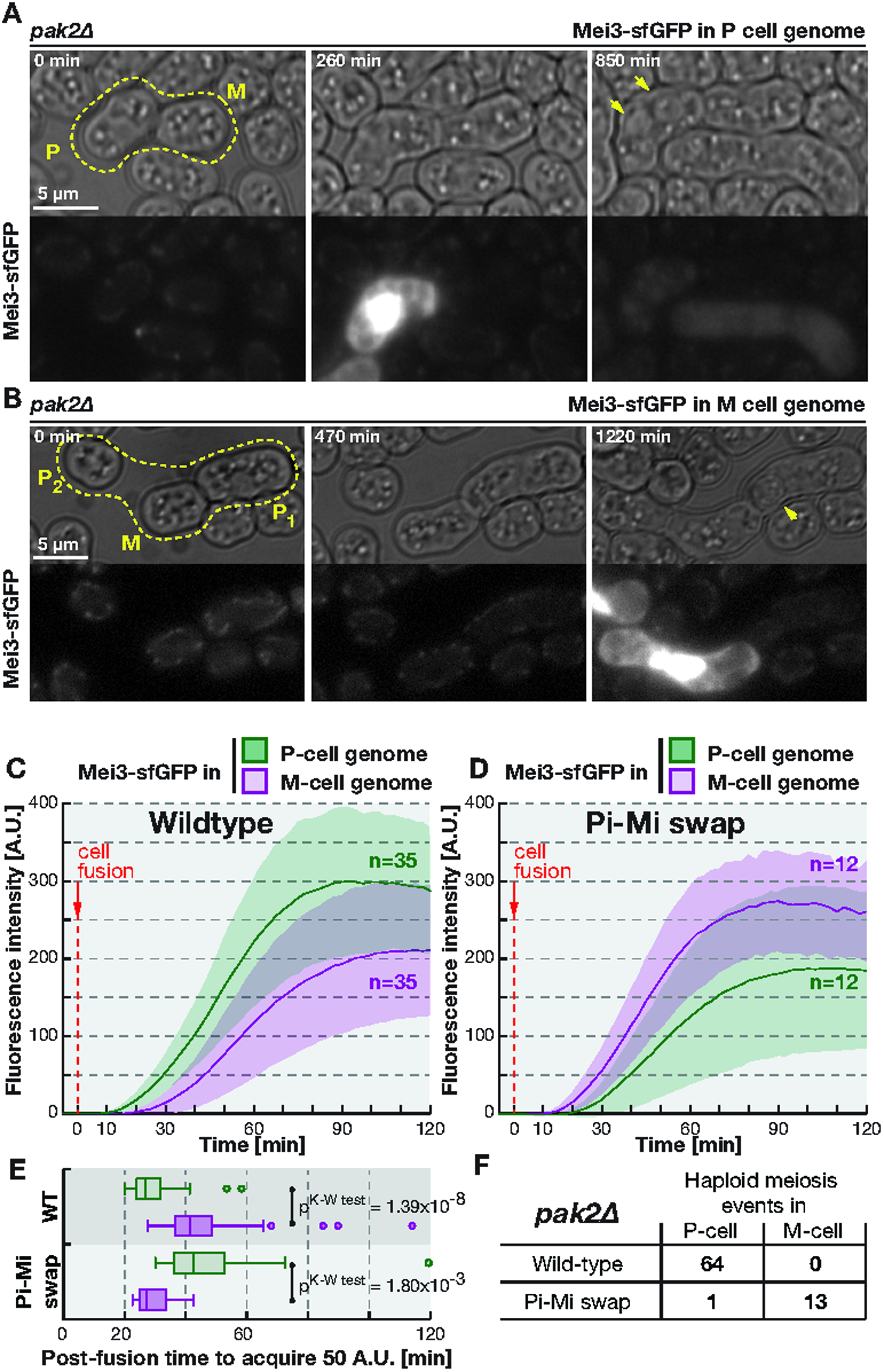
Meiotic inducer Mei3 is rapidly induced from the genome of the cell expressing homeodomain factor Pi. **(A)** Time-lapse showing expression of Mei3-sfGFP in the *pak2Δ* spore-forming P-cell (arrowhead) when *mei3* was fluorescently tagged only in the P-cell. Note signal absence in the M-cell. **(B)** Time-lapse showing transient cell fusion between *pak2Δ* M- and P1-cells, with the latter producing spores (arrowhead). Fluorescence tagging of *mei3* in the M-cell led to undetectable Mei3-sfGFP signal during transient fusion, but strong induction when M- and P2-cells completely fuse. **(C-D)** Quantification of fluorescence signal from zygotes of heterothallic wildtype (C) or cells with swapped Mi-Pi (D), where only one partner has Mei3 fused to sfGFP. Transfer of cytosolic mCherry between partners was used to observe cell fusion set to time zero. Solid lines represents average values and shaded areas include two standard deviations. **(E)** Box-and-whiskers plot (see Methods) of data shown in panels (C) and (D) with Kruskal-Wallis test p^value^ displayed. **(F)** Table reporting incidence of haploid meiosis in P- and M-cells for mating mixtures of *pak2Δ* cells with native or swapped expression of Pi and Mi.

Mei3 expression is under the regulation of two cell type-specific factors, the P-cell-specific homeodomain protein Pi and the M-cell-specific 42 amino acid peptide Mi^7^,^8^. To test whether these govern the asymmetric induction of *mei3* from parental genomes, we swapped the coding sequences of Pi and Mi. Without affecting mating and sporulation efficiencies nor spore viability (Fig. S2B-D), the Pi-Mi swap resulted in Mei3 being expressed first from the M-cell genome (Fig. 2D-E, S2E and Mov. S7), and transient fusion inducing haploid meiosis in M-cells (Fig. 2F, S2F-G and Mov. S8, 13 out of 14 instances). Thus, asymmetric zygotic transcription of *mei3* expression is governed by the cell type-specific expression of Mi and Pi, with *mei3* being induced first from the genome of the Pi-expressing cell.

## Pi and Mi nuclear-cytosolic shuttling underlies their asymmetric activity

To explore how parental genomes are differentially regulated by Pi and Mi, we tagged each with sfGFP at their native genomic loci. Mi-sfGFP expression commenced during early mating, producing a faint cytoplasmic signal in M-cells. Unexpectedly, after fusion, Mi-sfGFP rapidly accumulated in the P-cell nucleus and only ∼5-minutes later in the M-cell nucleus (Fig. 3A, S3A and Mov. S9). In *pak2*Δ cells undergoing transient cell fusion Mi-sfGFP transferred from the M- to the P-cell and accumulated in the nucleus of the P-cell, which then underwent haploid sporulation (Fig. 3B, n>10). Importantly, Mi did not accumulate in the nucleus of the M-cell that underwent transient cell fusion. We conclude that Mi nuclear accumulation requires a P-cell-specific factor that remains asymmetrically distributed upon transient cell fusion.

**Figure 3.**
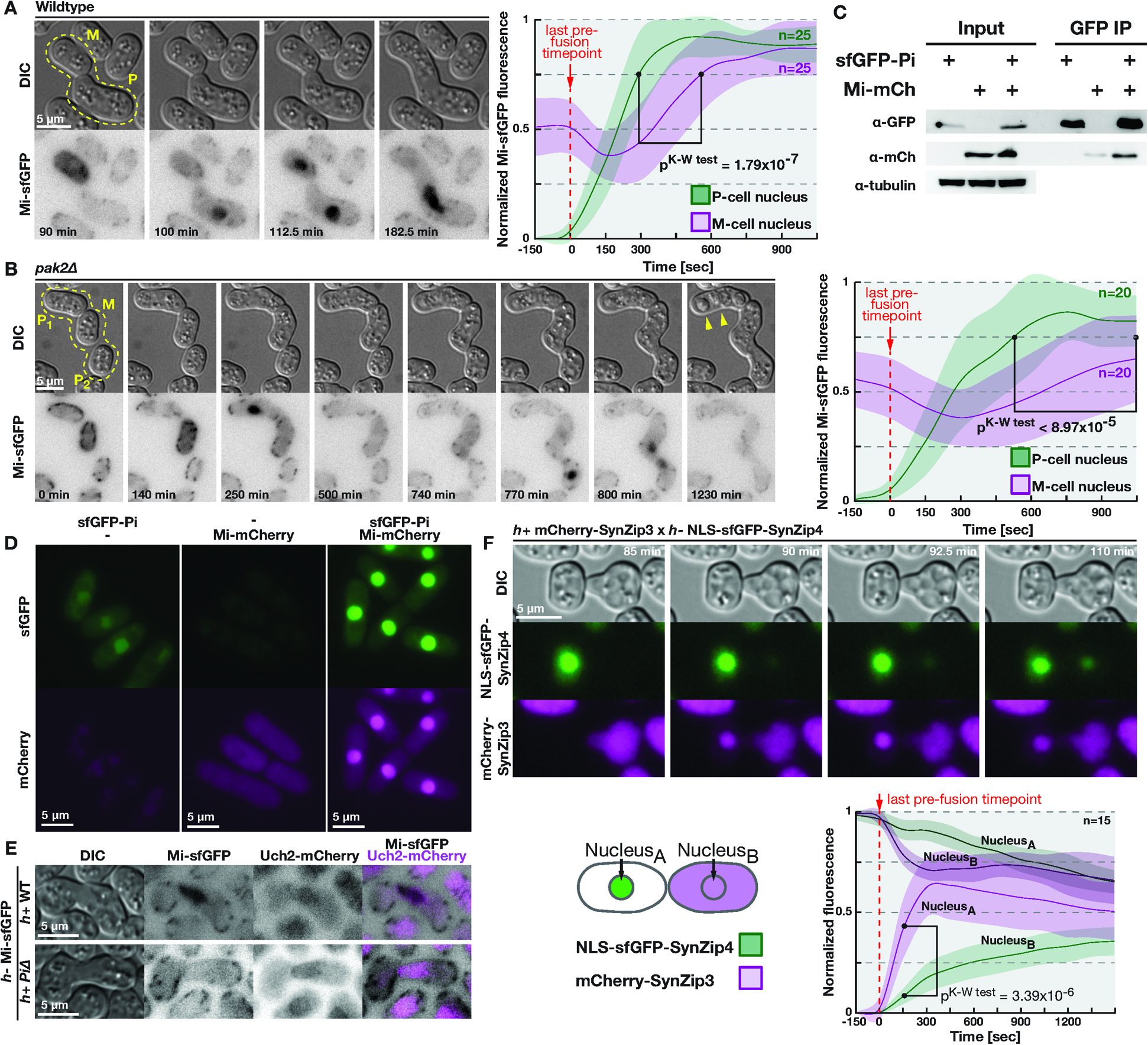
Meiotic regulators Pi and Mi asymmetry localize in early zygotes. **(A)** Left panel shows homothallic cells where M-cell-specific cytosolic Mi-sfGFP rapidly accumulates in the P-nucleus after partners fuse, and only subsequently in the M-nucleus. Note that the dotty signal at cell cortex in the green channel is background fluorescence, likely produced by mitochondria. Right panel quantifies the Mi-sfGFP signal in both partners’ nuclei with solid lines representing average values, shaded areas depicting two standard deviations and Kruskal-Wallis test p^value^. **(B)** Left panel follows transient cell fusion between homothallic *pak2Δ* partners, which resulted in exchange of Mi-sfGFP and its rapid accumulation in the P1-nucleus without any notable accumulation in the M-nucleus. Note that after M-cell switches partners, Mi also accumulated first in the P2-nucleus. Right panel quantification as in (A). **(C)** Cell lysates (Input lanes) obtained from interphase cells expressing sfGFP-Pi or Mi-mCherry or both were immunoprecipitated with GFP affinity beads and bound proteins (GFP IP lanes) analyzed by western blotting for each fluorophore and tubulin. Note co-immunoprecipitation of Mi-mCherry with sfGFP-Pi. **(D)** Micrographs of exponentially growing *mei3*Δ cells overexpressing, as indicated, sfGFP-Pi, Mi-mCherry or both proteins. Note nuclear localization of sfGFP-Pi and of Mi-mCherry only upon Pi co-expression. **(E)** Images of zygotes produced by M-cells expressing Mi-sfGFP with wildtype (top) and P-cells lacking Pi (bottom). Note that P-genome Pi is required for Mi accumulation in the nuclei labeled by Uch2-mCherry. **(F)** Top panel follows the mating of wildtype cells expressing NLS-sfGFP-SynZip4 and mCherry-SynZip3. Note rapid accumulation of the red fluorophore in the partner’s nucleus upon fusion and delayed exchange of the green fluorescence between partners. Bottom panel quantifies both fluorescent signals in the two nuclei, as labeled on the scheme, and is presented as in (B).

Several lines of evidence indicate that this P-cell-specific factor is Pi. First, Pi and Mi form complexes *in vivo* (Fig. 3C). Second, Pi was localized to the nucleus: N-terminal tagging of Pi with sfGFP produced a very low-intensity signal in nuclei in both P-cells and zygotes (Fig. S3B and Mov. S10). This nuclear localization was more evident in *fus1Δ* cells, which attempt, and fail, to fuse for a long time (Fig. S3C, Mov. S10), or upon overexpression in interphase cells (Fig. 3D). Third, Mi-mCherry overexpressed in interphase cells was cytosolic but accumulated in the nucleus when sfGFP-Pi was co-overexpressed (Fig. 3D). Finally, deletion of Pi prevented Mi-sfGFP nuclear enrichment after fusion (Fig. 3E, Mov. S11).

These data are consistent with a model (Fig. S3D) where Pi accumulates in the P-gamete nucleus while Mi is present throughout the nucleo-cytoplasm of the M-gamete. Upon cell fusion the 5kDa Mi is rapidly exchanged between partners and trapped by Pi in the P-nucleus where both proteins form a complex to rapidly induce Mei3 expression. Expression of Mei3 from the M-genome is postponed due to delayed exchange of Pi between partners.

We considered several reasons for the observed rapid, asymmetric nuclear localization of Mi. The 5kDa Mi peptide may diffuse significantly more rapidly than its 19kDa partner Pi^24^. However, tagging Mi with 27kDa sfGFP did not abrogate the asymmetry, as shown above, suggesting the asymmetry does not simply rely on protein sizes. The observed reversal of asymmetry upon Mi-Pi swap however indicates that the asymmetry does not depend on other cell type-specific factors. We thus considered whether distinct subcellular localization of Pi and Mi to the nucleus and the cytosol underlie the asymmetry. To mimic Pi and Mi characteristics, we constructed an artificial system consisting of synthetic hetero-specific short coiled-coils SynZip3 and SynZip4^25^, the former fused to mCherry and the latter to sfGFP and a nuclear localization signal (NLS), each expressed in one partner. Remarkably, upon cell-cell fusion, the cytosolic marker accumulated in the partner’s nucleus immediately, whereas the equilibration of the nuclear marker was delayed, thus mimicking the behavior of Mi and Pi (Fig. 3F, Mov. S12). The observed asymmetry was even more striking in instances of transient cell fusion (Fig. S3E, Mov. S12). When both proteins lacked an NLS (Fig. S3F, Mov. S12), asymmetric localization was not observed upon fusion. We conclude that the observed asymmetry is inherent to the distinct sub-cellular localization of Mi and Pi, a design that promotes a more rapid Mei3 transcription from the P-genome.

## Rapid induction of Mei3 transcription prevents re-fertilization and polyploid formation

To address the physiological function of this fast Mei3 induction, we delayed Mei3 expression in two distinct ways. First, we placed Mi in M-cells under control of a P-cell specific *p*^*map3*^ promoter, such that Mi (and consequently Mei3) becomes expressed only after successful fusion, once P-cell-specific transcription factors activate the *p*^*map3*^ promoter in the M-genome. This delayed Mei3-sfGFP expression by ∼30 minutes (Fig. 4A, S4A and Mov. S13). Second, we simply deleted *mei3* from the P-genome, such that Mei3 is expressed only from the M-genome, and thus delayed by ∼15 minutes (Fig. 2E). While neither approach affected mating or sporulation efficiencies upon complete fusions of otherwise wildtype cells (Fig. S4B, n=3×300), both methods fully prevented haploid sporulation upon transient fusion of *pak2* mutants. Instead, we observed persistent mating behaviors with both partners repeatedly attempting to form a zygote (note the post-fusion growth of mating projections in Fig. 4B, 4C and Mov. S14, n=10). Deleting *mei3* in *pak2*Δ M-cells, however, did not affect haploid sporulation in the P-cell (Fig. S4C and Mov. S14, n>10).

**Figure 4.**
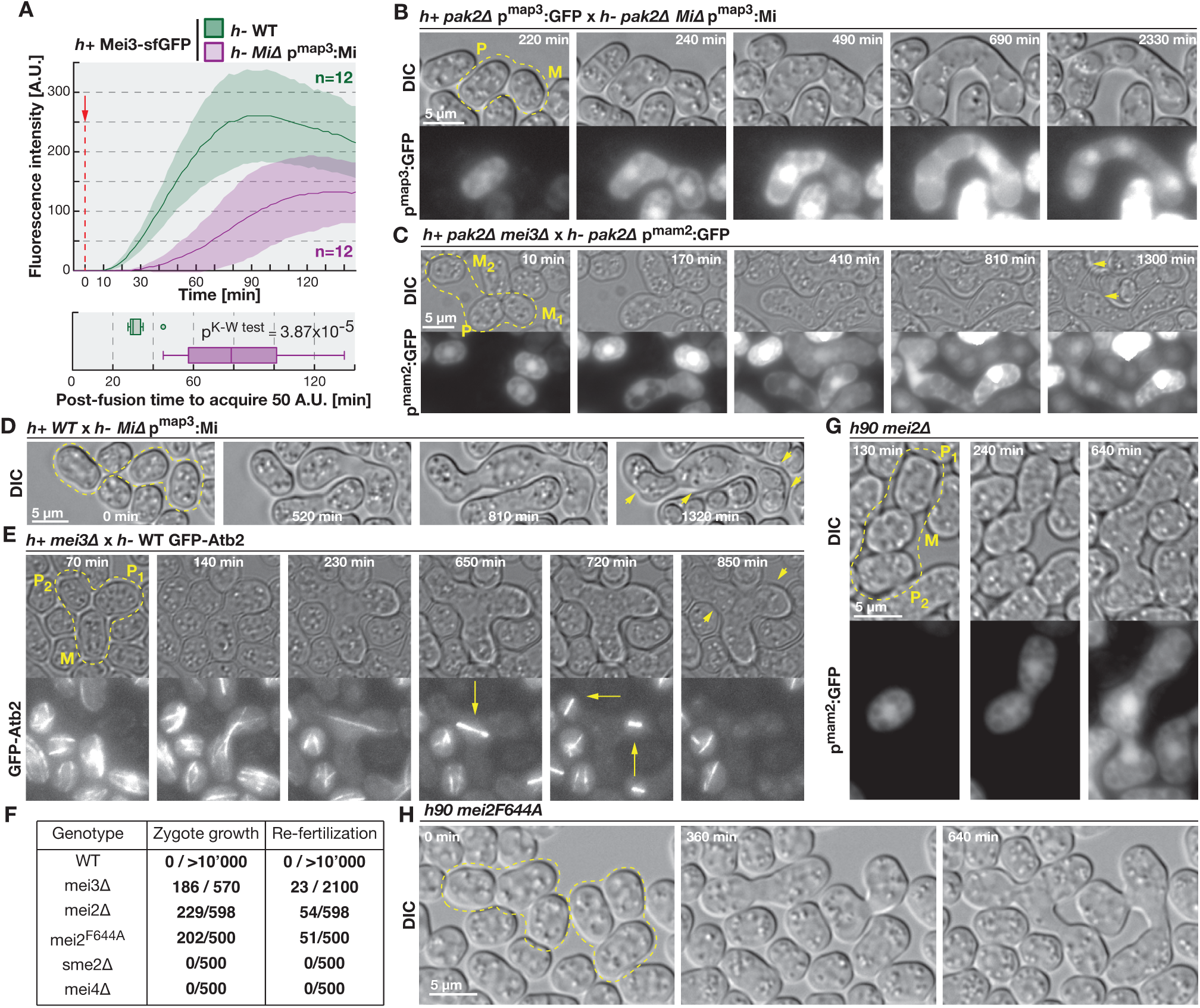
Rapid induction of Mei3-Mei2 signaling is required to suppress mating in zygotes and prevent polyploid formation. **(A)** Quantification of P-genome encoded Mei3-sfGFP fluorescence in zygotes from either wildtype M-cells (green) or M-cells with Mi expression delayed post-fusion due to control by P-cell specific, *p*^*map3*^, promoter (magenta). Data is presented as in Fig. 2C and 2E. **(B)** Time-lapse showing mating of *pak2Δ* P-cells expressing cytosolic GFP and M-cells where Mi expression is under the *p*^*map3*^ promoter. Note that transient fusion, observed as initial GFP exchange followed by signal build up in only one partner, did not induce meiosis in either partner. Instead, partners continue to extend mating projections until complete fusion occurs and only then make spores. **(C)** Time-lapse showing mating of *pak2Δ* mutant cells where *mei3* has been deleted from the P-partner and cytosolic GFP is expressed in M-cells. Transient cytosolic exchange between P and M1 cells, failed to induce sporulation and instead partners continue mating with each other until the P-cell eventually switches partners. **(D)** Time-lapse showing a zygote fusing with an additional partner upon mating of wildtype P-cells with M-cells expressing Mi under regulation of the *p*^*map3*^ promoter. Note formation of spores in the now triploid zygote. **(E)** Time-lapse showing crosses where *mei3* was specifically deleted in P-cells and M-cells expressed GFP-α-tubulin. Note fusion of three partners, formation of meiotic spindles (arrows) and spores (arrowheads). **(F)** Table showing incidence of re-fertilization and zygotic growth in mating mixtures of homothallic strains with indicated genotypes. **(G)** Time-lapse showing repeated fertilization of homothallic *mei2Δ* cells, marked by exchange of cytosolic GFP produced by M-partner. **(H)** Time-lapse of homothallic *mei2*^*F644A*^ cells carrying a point mutation in the RNA recognition motif. Note the formation of triploid zygotes.

Remarkably, when Mei3 expression was delayed by either method in otherwise wildtype crosses, zygotes formed but then continued to engage additional partners (Fig. 4D and Mov. S15-16, n=18 for Mi expression delay and 18 for *mei3* deletion in P-cells, out of 7’000 events each). Multi-partner zygotes entered meiosis, as shown by microtubule labeling, and formed four, likely aneuploid spores (Fig. 4E, S4E, Mov. S15-16). By contrast, we rarely observed triploid zygotes when *mei3* was deleted from the M-cell genome (2 of n > 7’000 events). Polyploid zygotes were never observed in wildtype (n > 10’000). In sum our data reveal that the rapid onset of Mei3 expression from the zygotic P-cell genome functions to prevent formation of polyploid zygotes and consequently aneuploid spores.

Furthermore, approximately 1% of all zygotes formed by homothallic *Pi*Δ*, Mi*Δ or *mei3*Δ strains not only did not enter meiosis, but were seen fusing with an additional partner (Fig. S4G-I and Mov. S17, n>500). A much larger fraction (>30%) of zygotes exhibited growth suggestive of mating behavior (Fig. 4F, Mov. S17-18). The frequency of re-fertilization events appeared to increase further with time but could not be quantified due to deterioration of imaging chambers.

Mei3 alleviates repression of the master meiotic regulator, the RNA binding protein Mei2^10^,^26^,^27^, which, in complex with *meiRNA*/*sme2*^9^, promotes meiosis by indirectly allowing expression of downstream factors including the forkhead transcription factor Mei4^28^,^29^. Both *mei4Δ* and *sme2*Δ zygotes arrested prior to spore formation but were never observed engaging additional partners (Fig. S4J-K). Conversely, nearly 10% of *mei2*Δ zygotes fused with additional partners (Fig. 4G and Mov. S18). Since a point mutation in Mei2 C-terminal RNA-recognition motif^9^ also resulted in ∼10% of multi-partner zygotes (Fig. 4H and Mov. S18), we conclude that the re-fusion block is reliant on Mei2 signaling, likely through RNA binding, but not in complex with *meiRNA*. These data establish that repression of mating in zygotes shares components with, but bifurcates from, meiotic induction.

Mechanisms preventing re-fertilization are present in evolutionarily divergent phyla to ensure ploidy maintenance across generations. Almost universally, these mechanisms involve rapid changes, be it through release of cortical granules^1^ and shedding of cell-surface receptors in mammals^2^, membrane depolarization in amphibians^1^,^30^, or degradation of pollen guidance cues in plants^31^. Our results now provide a first glimpse into the mechanism blocking re-fertilization in fungi, by showing that rapid activation of zygotic transcription promotes discontinuation of mating in yeast zygotes. The bi-partite design of the Mi-Pi transcription factor, reminiscent of that of hormone nuclear receptors and their ligands^32^, favors post-fusion activation speed, which is limited only by the rate at which the Mi peptide reaches the already nuclear-localized homeodomain Pi protein. Because this simple two-component system inherently leads to asymmetric zygotic expression, asymmetry may naturally follow from the selective pressure to rapidly block re-fertilization. The observation that even a transient cytoplasmic connection is sufficient for formation of an active complex raises the possibility that such transcription factor design may also be used to impart cell fate changes upon, or build sensors to monitor, other instances of transient cytoplasmic bridge formation.

## Materials and Methods

Detailed materials and methods are provided in a supplemental file appended below.

## Acknowledgements

We thank Benoit Arcangioli and Geneviève Thon for strains and advice, Magdalena Marek for help with flow cytometry, Felipe Bendezú and Serge Pelet for strains and reagents, and Richard Benton, Tonni Grube Andersen, Stefan Gruber, Sara Mitri, Jan-Willem Veening and Martin lab members for critical reading of the manuscript. This work was supported by an EMBO long-term fellowship to AV and an ERC Consolidator Grant (CellFusion) and a Swiss National Science foundation grant (31003A_155944) to SGM.

## Supplemental Movie legends

**Movie 01. *pak2Δ* cells undergo haploid meiosis**

Time-lapse of homothallic wildtype (top panel) and *pak2*Δ (bottom panel) cells expressing GFP-α-tubulin in green and nuclear marker Uch2-mCherry and spindle pole body marker Pcp1-mCherry in magenta. Note that in the outlined mating pairs of *pak2Δ* cells, one partner forms meiotic spindles (arrows at time points 5:40h to 6:30h) while the other partner maintains its nucleus and interphase microtubule organization. Aberrant type IIIa and IIIb asci are indicated in the last time point.

**Movie 02. *pak2Δ* cells undergo transient cell fusion followed by sporulation in the P-cell**

In the wildtype (left panel) or *pak2Δ* mutant (right panel) background, we mated *h-* cells expressing Myo52-GFP with *h+* cells expressing Myo52-tdTomato and GFP from the P-cell specific *p*^*map3*^ promoter. GFP and Myo52-GFP are depicted in green and Myo52-tdTomato in magenta. Cell fusion is observed as exchange of cytosolic GFP between partners. Note that transient cell fusion between *pak2*Δ M- and P_1_-cells is followed by sealing of the fusion pore and sporulation in the P-cell, while the M-cell engages another partner.

**Movie 03. Transient cell fusion between *pak2Δ* cells leads to sporulation in the P-cell.**

Shown are homothallic *pak2Δ* cells expressing GFP from the M-cell specific p^mam2^ promoter. Note that in the indicated mating pairs cytosolic GFP exchange is followed by fusion pore closure and build up of the GFP signal only in M-cells while indicated P-cells proceed to sporulate.

**Movie 04. Transient cell fusion between *fus1Δ* and wildtype cells leads to sporulation in the P-cell.**

Time-lapse imaging showing that mating between wildtype and *fus1*Δ heterothallic strains with indicated genotypes results in instances of transient cell fusion (outlined cell pairs) followed by haploid sporulation in the P-cell.

**Movie 05. Transient cell fusion induces Mei3 expression specifically in the P-cell.**

Time-lapse of mating between heterothallic *pak2*Δ strains for which *mei3* is fused to sfGFP in only one partner, as indicated. Note that upon transient cell fusion between outlined mating pairs fluorescent signal is observed only if the P-cell codes for Mei3-sfGFP (left panel). When Mei3 was fluorescently tagged in M-cells only (right panel) no detectable fluorescent signal was observed in pairs undergoing transient fusion, which produced spores in the P-partner (arrowheads). Note that Mei3-sfGFP is produced upon complete fusion whether it is encoded in one or the other partner.

**Movie 06. Mei3 is first induced from the P-cell genome of wildtype zygotes.**

Time-lapse showing mating of wildtype M-cells with P-cells expressing cytosolic mCherry and, as indicated, Mei3 tagged with sfGFP in either the P-partner (top, examples 1-4) or in the M-partner (bottom, examples 5-8). Cell fusion, visualized as exchange of cytosolic mCherry, is set as time zero in all examples. Note more rapid induction of the green fluorescence when *mei3-sfGFP* is encoded in the P-genome.

**Movie 07. Mei3 is preferentially induced from the M-cell genome of zygotes produced by partners with swapped expression of Pi and Mi.**

Shown is mating of heterothallic strains with swapped Pi and Mi expression where P-cells express cytosolic mCherry and, as indicated, Mei3 tagged with sfGFP in either the P-partner (top, examples 1-3) or M-partner (bottom, examples 4-6). Cell fusion, visualized as exchange of cytosolic mCherry, is set as time zero in all examples. Note more rapid induction of the green fluorescence when *mei3-sfGFP* is encoded in the M-genome.

**Movie 08. Transient cell fusion between cells with swapped expression of Pi and Mi leads to sporulation in the M-cell.**

Time-lapse showing that transient fusion between cells with swapped Pi and Mi expression induced sporulation in the M-cell. The transient fusion, evident as exchange of cytosolic mCherry expressed from the *p*^*map3*^ promoter in outlined pairs, was caused by either *pak2* deletion in both partners (left panel) or *fus1* deletion in only one partner (right panel). Note that the M-cells experiencing transient fusion eventually sporulate (arrowheads), while P-cells engage proceed to engage additional partners.

**Movie 09. Post-fertilization Mi first accumulates in the P-cell nucleus.**

Time-lapse imaging shows Mi-sfGFP localization during mating of wildtype (top and middle panels) and *pak2Δ* cells (bottom panel). The cells in the middle panel also express the nuclear marker Uch2-mCherry (magenta). The faint cytosolic Mi-sfGFP signal initially present in the M-cell rapidly accumulates in the P-cell nucleus post-fusion. Note that the dotty signal at cell cortex in the green channel is background fluorescence, likely produced by mitochondria. The asymmetric Mi nuclear localization is better observed with higher temporal resolution of imaging in the top panel. 10-minute imaging interval in the middle panel captures asymmetric Mi-sfGFP localization in only some examples. Transient cell fusion (bottom panel) is associated with Mi enrichment only in the nucleus of the P_1_-cell that eventually sporulates (arrowheads), and the asymmetric Mi-sfGFP nuclear accumulation is also evident in M-cell fusion upon complete fusion with the P_2_-partner.

**Movie 10. Pi localizes to the nucleus pre-fertilization.**

Time-lapse shows sfGFP-Pi localization during mating of homothallic wildtype (top) and *fus1Δ* cells (bottom) also expressing the nuclear marker Uch2-mCherry (magenta). Note the faint nuclear signal of Pi (arrows) that exhibits movements that mirror that of the Uch2-labelled nucleus. The dotty signal in the green channel at cell cortex is a background signal that can be observed in all cells.

**Movie 11. Nuclear Mi recruitment is Pi-dependent.**

Time-lapse imaging shows Mi-sfGFP expressing M-cells mating with either wildtype (top) or *Pi*Δ (bottom) P-cells. Note that the accumulation of Mi to the nuclei, visualized by Uch2-mCherry (magenta), is abolished in zygotes lacking Pi. 10-minute imaging interval captures asymmetric Mi-sfGFP localization in only some examples.

**Movie 12. Asymmetric nuclear accumulation of Pi-Mi complex upon fertilization can be mimicked with a pair of synthetic peptides.**

Time-lapse imaging shows mating of heterothallic wildtype (left and right panels) or *pak2Δ* cells (middle panel) expressing either mCherry (magenta) or sfGFP (green) tagged with indicated SynZip peptides that promote formation of fluorophore hetero-dimers upon fertilization. In the left and middle panel the sfGFP localizes to the nucleus due to an N-terminal nuclear localization signal (NLS) which is absent from the construct presented in the right panel. Cytosolic mCherry rapidly re-localizes to the nucleus of the NLS-sfGFP-expressing cell upon cell fusion (left and middle panels), but not the sfGFP-expressing cell (right panel). By contrast, the nuclear, but not cytosolic, sfGFP construct is slow to exchange between zygotic nuclei. Upon transient cell fusion in *pak2*Δ cells (middle panel), evident from spore formation in the P-cell (arrowheads), the asymmetric nuclear localization of the fluorophores is persistent. Note that in addition to the cytosolic signal, the mCherry-SynZip3 construct produced background vacuolar fluorescence, likely due to pre-fusion degradation.

**Movie 13. Placing Mi-expression under P-cell specific promoter delays induction of Mei3 in zygotes**

Time-lapse showing mating of wildtype P-cells encoding Mei3-sfGFP and cytosolic mCherry with either wildtype M-cells (top, examples 1-3) or M-cells where Mi-expression is under the regulation of the P-cell specific *p*^*map3*^ promoter (bottom, examples 4-6). Cell fusion, visualized as exchange of cytosolic mCherry, is set as time zero in all examples. Note the retarded induction in the mutant as compared to wildtype.

**Movie 14. Delayed induction of Mei3 prevents haploid meiosis in *pak2Δ* cells that undergo transient cell fusion.**

Time-lapse imaging shows mating of *pak2*Δ mutant cells where Mei3 expression was delayed by either placing Mi under the P-cell specific *p*^*map3*^ promoter (left panel) or by deleting *mei3* specifically in the P-cell (middle panel). Note lack of sporulation in the P-partner that underwent transient cell fusion, observed as exchange of cytosolic GFP followed by signal increase only in the M-partner. Instead, partners continue to grow mating projections until they fuse (left panel) or engage another partner (middle panel). The right panel shows that deletion of *mei3* in the *pak2*Δ M-partner did not affect occurrence of haploid meiosis (arrowheads) in the *pak2*Δ P-cell upon transient cell fusion.

**Movie 15. Delayed induction of Mi leads to formation of polyploids.**

Time-lapse imaging shows mating of wildtype P-cells with M-cells that have Mi expression under the P-cell specific *p*^*map3*^ promoter. Note the fusion between the diploid zygote and haploid cell visualized as cytosolic mCherry exchange (right panel) and formation of spores (arrowheads) by the triploid zygote (left panel).

**Movie 16. Delayed induction of mei3 results in re-fertilization.**

Time-lapse imaging shows mating of *mei3Δ* mutant P-cells with wildtype M-cells (left panel) or M-cells with fluorescently labeled tubulin (middle panel) or cytosol (right panel). Note formation of polyploid zygotes between outlined cells that proceed to form meiotic spindles (middle panel, arrows) and sporulate (arrowheads)

**Movie 17. Zygotes lacking Pi, Mi and Mei3 fail to repress mating and form polyploids.** Time-lapse imaging shows mating of homothallic strains lacking indicated genes. Note consecutive fusion between three partners (*Pi*Δ panels) as well as fusion of two zygotes (*mei3Δ* and *MiΔ* panels). In the left and right panels, P-cell-specific and M-cell-specific fluorescence, respectively, is used to highlight fusion events. The arrows in the middle panel point to instances of zygotes forming mating projections.

**Movie 18. Zygotes lacking *mei2* or carrying a mutation in its second RNA-recognition motif fail to repress mating.**

Time-lapse shows mating of homothallic *mei2Δ* cells expressing cytosolic GFP (left panels) or *mei2*^*F644A*^ (right panels) mutant cells. Note that outlined zygotes fail to repress mating and proceed to make polyploids.

## Materials and Methods

### Growth conditions

The growth conditions used for experiments are detailed in ref.^1^ and the overview of experimental procedure is presented in Fig. S5A. Briefly, freshly streaked cells were inoculated into MSL+N media^2^ and incubated overnight at 25°C with 200 rpm shaking to exponential phase. The following evening cultures were diluted to O.D._600nm_=0.025 in 20 ml of MSL+N media and incubated overnight at 30°C with 200 rpm agitation to exponential phase. Experiments on exponentially growing cultures were performed at this point. For time-lapse imaging of mating, homothallic cells or 1:1 mixtures of heterothallic cells were pelleted for one minute at 1000g in microcentrifuge tubes and washed three times in 1ml of MSL-N media^2^. Cells were then diluted in 3ml of MSL-N media to final O.D._600nm_=1.5 and incubated at 30°C with 200 rpm agitation for 4-6 hours. Finally cells were mounted onto MSL-N agarose pads, covered with a coverslip and the chamber was sealed using VALAP (Vaseline, Lanolin, Paraffin; 1:1:1).

For flow cytometry and **quantifications of mating and sporulation efficiencies**, ∼3×10^7^ of MSL-N washed cells were re-suspended in 20 µl of MSL-N media and spotted onto MSL-N 2% agar plates and incubated at 25°C for the indicated number of hours. The number of unmated cells, unsporulated zygotes and sporulated zygotes was determined using transmitted light microscopy, and mating and sporulation efficiencies were calculated using the following formulas:

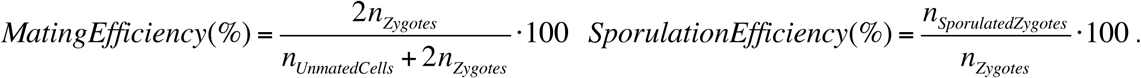

The reported results are from three replicates, error bars denote standard deviation and p-values were calculated using Welch’s *t*-test.

### Flow cytometry

We collected the mated cell suspensions from MSL-N plates and resuspended samples in 1ml of MSL-N media containing 10µl of glusulase (NEE154001EA, Perkin-Elmer, Waltham, USA). After overnight incubation at 30°C we visually verified that all cells except spores had lysed upon glusulase treatment. The samples were washed three times with MSL-N and resuspended in 3ml of water containing 2-5 µg/ml of Hoechst 33342 DNA dye (B2261, Sigma-Aldrich, St. Louis, USA) and incubated for 15-30 minutes at room temperature. Samples were immediately analyzed on Becket-Dickson Fortessa with proprietor software platform using the 355nm laser line with 450/50 filter. The experiment was run in duplicate.

### Strains

All strains are reported in Table S1. Standard fission yeast genetic and molecular biology tools were used in the study^3^. Sequences of mutants obtained in this study are provided as annotated, GenBank-formatted sequences (see Supplemental Sequences).

The **mating loci** of strains used in the study are illustrated in Fig. S5B. Mi and Pi deletions remove the ORF of the genes. sfGFP tagging of Pi and Mi was done at the N- and C-terminus respectively.

In homothallic strains, the ***mat1* locus** determines the cell mating type and switches between sequences encoded by the silent *mat2* and *mat3* loci [reviewed in ref.^4^]. Importantly, a H1Δ17 mutation at the *mat1* locus H1-homolgy box prevents mating type switching^5^. As depicted in the Fig. S5B, in the synthetic *mat1* locus sequences, the wild-type H1-homology box was replaced with the one amplified from the PB9 strain carrying the H1Δ17 mutation (a kind gift from Benoit Arcangioli, Institut Pasteur, Paris, France). The recombinant DNA also contain a selection cassette and homology arms that targets the constructs to the *mat1* locus of homothallic cells. The described sequences were cloned and amplified inside the pBlueScript plasmid. Finally the indicated restriction enzymes (see Supp.Seq.1-4) were used to extract the fragment of interest and transform it into *h90* wild type cells. Correct integration was verified by PCR and/or inability to perform mating type switching.

***mat2* and *mat3* loci** that reside in heterochromatic DNA and thus cannot be targeted by homologous recombination in wild-type cells were manipulated as described in ref.^6^. An overview of the experimental procedure is presented in Fig. S6A. The *mat2* locus manipulations were performed in the TP1 strain (a kind gift from Genevieve Thon, University of Copenhagen, Copenhagen, Denmark), which lacks the *clr3* gene, to allow recombination at the *mat2* locus. Furthermore, the TP1 strain carries *ade6+* and *ura4+* cassettes in the vicinity of the *mat2* locus. The TP1 strain was transformed with DNA fragments carrying the designed *mat2* locus and flanking wild-type sequences. 5-FOA resistant, adenine auxotrophs were selected and tested for correct integration by PCR. The *mat2* mutant locus was then introduced in otherwise wild-type cells via genetic crosses. *mat3* manipulations were similarly performed in the PG3089 strain (a kind gift from Genevieve Thon), which lacks the *clr4* gene and carries the *ura4+* cassette in the proximity of the *mat3* locus. The sequences of the mutant *mat2* and *mat3* loci are provided (see Supp.Seq.5-8).

For **Pi overexpression** we used the constitutive *tdh1* promoter comprised of 1000bp upstream of the *tdh1* start codon to drive expression of Pi N-terminally tagged with sfGFP. This construct was cloned into plasmids that contain the budding yeast ADH1 transcriptional terminator sequence, BleMX selection cassette and sequences targeting integration at either *ura4* or *his5* locus (Supp.Seq.9-10). AfeI digested *ura4* targeting plasmid and StuI digested *his5* targeting plasmid were transformed in *ura4* and *his5* mutant cells respectively. Prototrophic, zeocin-resistant clones were selected and checked for correct integration by PCR and genetic crosses.

For **Mi overexpression** we used either the *tdh1* or the *nmt41* promoter to drive expression of Mi that was C-terminally tagged with mCherry. These constructs were cloned into plasmids that target either *ura4* or *lys3* locus and contain kanMX or natMX selection cassettes respectively (Supp.Seq.11-12). AfeI digested *ura4* targeting plasmid and BstZ17I digested *lys3* targeting plasmid were transformed in corresponding auxotrophic strains and prototrophic clones with adequate antibiotic resistance were selected and checked for correct integration by PCR and genetic crosses.

*mei3Δ* mutant background was used to prevent induction of meiosis by **co-expression of Mi and Pi**^7^,^8^ as was the case for control strains expressing individual proteins.

For **Mi expression from the P-cell-specific *p*^*map3*^ promoter** we used 2063bp upstream of the *map3* start codon followed by the Mi ORF sequence, the budding yeast ADH1 transcriptional terminator sequence and BleMX selection. This construct was cloned inside the pJK148 plasmid^9^ that targets the construct to the *leu1* locus after NdeI digestion (Supp.Seq.13). The linearized plasmid was transformed into *h*-cells lacking Mi and a G418 resistant transformant that efficiently sporulated when crossed with wildtype *h+* cells was selected. The same transformation clone was used to introduce the *pak2Δ::hphMX* mutation as described below.

***pak2Δ::ura4+*** mutant strain was derived from ref^10^. For the ***pak2Δ::hphMX*** we cloned the 430bp of 3’UTR^pak2^ followed by 399bp of the 5’UTR^pak2^ into the pFA6a-hphMX vector which created an AfeI site between the two UTR’s. The obtained plasmid (Supp.Seq.14) was linearized with AfeI, transformed into cells and hygromycin-B resistant colonies were selected and tested for correct integration by PCR.

***mei2Δ::kanMX*** and ***mei3Δ::kanMX*** mutant strains were derived from the *S. pombe* deletion library strains S7B02 and S20A08 respectively (Bioneer, Daejeon, Republic of Korea) and were verified by PCR.

***sme2Δ::ura4+*** and ***mei4Δ::ura4+*** mutations were derived from Japan National BioResource Project strains FY7237 and FY7361, respectively.

***fus1Δ::natMX*** was generated using methodology detailed in ref.^11^. Briefly, primers with 80bp homology to sequences immediately flanking fus1 ORF (see Table S2) were used to PCR amplify the natMX selection cassette from the pFA6a-natMX plasmid and transformed into cells. Nourseothricin resistant clones were selected and genotyped by PCR.

For construction of ***mei2* point mutant** we first cloned the 3’ region (sequence from 313-672bp downstream *mei2* stop codon) followed by the *mei2* gene (sequences from 487bp upstream of the start codon to 312bp downstream of the stop codon), which creates an AfeI restriction enzyme between the two fragments. This construct was inserted in the pFA6a-hphMX plasmid and we introduced the F644A mutation by site-directed mutagenesis (Supp.Seq.15-16). Linearization of the plasmid with AfeI targets the construct to replace the genomic *mei2* gene upon transformation. The hygromycin-B resistant clones were selected and the correct integration replacing the wild-type *mei2* was confirmed by PCR.

The strains with a **fluorescently labeled genomic locus**^12^ were derived from FY13708 strain obtained from the Japan National BioResource Project. In these cells the *lys1+* locus is genetically linked to the *LacO* sequence array that binds the GFP-LacI expressed from the *dis1* promoter at the *his7+* locus.

For **microtubule visualization** we used cells with SV40 promoter driven expression of GFP-α-tubulin^13^ derived from a FC1234 strain, a kind gift from Fred Chang’s group.

For ***mei3* fluorophore tagging**, yeast codon optimized sfGFP (a kind gift from Michael Knop, Heidelberg University, Germany) was introduced into a pFA6a-natMX vector to obtain the pFA6a-sfGFP-natMX vector (Supp.Seq.17). The plasmid was then used as a template for PCR with primers that amplify sfGFP followed by natMX cassette and carry 80bp homology to sequences immediately flanking *mei3* STOP codon (see Table S2). The PCR product was transformed into wild-type cells, and nourseothricin resistant clones were selected and genotyped by PCR. All mei3-sfGFP expressing strains used in the study were derived from the same transformation clone by genetic crosses.

For C-terminal mCherry **tagging of nuclear marker *uch2* and spindle pole body marker *pcp1***, we used the pFA6a-mCherry-natMX and pFA6a-mCherry-kanMX plasmids respectively. In both cases, we cloned the 3’UTR region of the gene followed by C-terminal fragment of the ORF keeping it in frame with the fluorophore (Supp.Seq.18-19). The obtained plasmids to tag *uch2* and *pcp1* were then treated with AfeI and AfeI+SnaBI restriction enzymes respectively, and the digested DNA was transformed into cells. Clones resistant to either nourseothricin or G418 were selected and genotyped by PCR.

**myo52-GFP** and **myo52-tdTomato** were introduced from strains FC857^14^ and ySM740 ^15^ respectively.

The construct to express the green and red fluorophores from the P-cell-specific *map3* promoter were previously described in ref.^16^. For the expression of **GFP from the M-cell-specific *mam2* promoter** we cloned the 438bp of promoter upstream of the start site into a pRIP-GFP plasmid that contains the *ura4+* selection cassette (Supp.Seq.20). Transformation of the *ura4*-mutant cells with the PmeI digested final plasmid targeted the construct to the native *mam2* promoter and conferred growth in absence of uracil.

Sequences encoding small interacting SynZip peptides^17^ were a kind gift from Serge Pelet, University of Lausanne, Switzerland. The SynZip3 and SynZip4 peptides were cloned at the C-terminus of mCherry and sfGFP respectively. Expression of mCherry-SynZip3 was driven from the *p*^*act1*^ promoter consisting of 822bp upstream of the *act1* start codon. The sfGFP-SynZip4 was driven from the *p*^*tdh1\**^ promoter comprised of 896bp upstream of the *tdh1* start codon. For the NLS-sfGFP-SynZip4 we introduced the SV40 nuclear localization signal at the sfGFP N-terminus and used the *p*^*tdh1*\*^ promoter comprised of 896bp upstream of the *tdh1* start codon. All constructs were integrated in the plasmid that carries the *ura4+* selection cassette and targets the *ura4* locus. All constructs are detailed in Supp.Seq.21-23. The AfeI linearized final plasmids were transformed into *ura4* mutants and uracil prototrophs were selected for further experiments.

### Co-immunoprecipitation

We grew 50ml of *mei3Δ* mutant cell cultures overexpressing Mi-mCherry or sfGFP-Pi or both to exponential phase and collected cells by centrifugation for one minute at 1000 rcf. From this point the experiment was performed on ice. Cells were transferred to microcentrifuge tubes, washed twice with ice-cold CXS buffer (50mM HEPES,pH7.5; 20mM KCl; 2mM EDTA; 1mM MgCl_2_; protease inhibitors) and re-suspended in 300µl of the CXS buffer and ∼200µl of zirconium beads were added. Cells were lysed using the bead beater with 10 cycles of 20-second beating and 40 seconds cooling on ice. The beads were removed and samples transferred to a new tube where the cell debris was pelleted by centrifugation for 15 minutes at 13’000g. We collected the supernatant and determined the protein concentration using the Bradford assay. We adjusted the samples to 1mg of total protein in 500µl of CXS buffer and set aside 30µl for western blotting analysis (Fig. 3A labeled as INPUT). The samples were then incubated with 30µl of GFP-Trap beads (gtma-20, Chromotek, Planegg-Martinsried, Germany) prewashed in CXS buffer. After one hour incubation on a rotating stand at 4°C beads were washed three times in CXS buffer and re-suspended in Laemmli buffer. Samples were analyzed with standard western blotting protocol. For primary antibodies we used 1:3’000 dilution of anti-GFP antibody (Cat.No. 11814460001, Roche, Basel, Switzerland) anti-RFP antibody (6G6, Chromotek, Planegg-Martinsried, Germany) and TAT-1 antibody (a kind gift from Keith Gull, University of Oxford, UK). We used the infrared fluorophore coupled secondary antibodies at 1:5’00 dilution (R-05061 and R05054, Advansta, Menlo Park, USA) and visualized the blots on the Fusion FX (Vilber, Collégien, France). The experiment was reproduced in three independent replicates.

### Microscopy and image analysis

To acquire images in Fig. 3D and S1C we used a spinning-disk confocal system that uses an inverted microscope (DMI6000B; Leica) equipped with an HCX Plan Apochromat 100x/1.46 NA oil objective and an UltraVIEW system (PerkinElmer; including a real-time confocal scanning head [CSU22; Yokagawa Electric Corporation], solid-state laser lines, and an electron-multiplying charge coupled device camera [C9100; Hamamatsu Photonics]). Stacks of z-series confocal sections were acquired at 0.3-µm intervals using Volocity software (PerkinElmer).

All other micrographs were obtained by wide-field microscopy performed on a DeltaVision platform (Applied Precision) composed of a customized inverted microscope (IX-71; Olympus), a UPlan Apochromat 100x/1.4 NA oil objective, a camera (CoolSNAP HQ2; Photometrics), and a color combined unit illuminator (Insight SSI 7; Social Science Insights). Images were acquired using softWoRx v4.1.2 software (Applied Precision). Principally we imaged a single Z-section with the exception of data presented in Fig. 1B and S1B where 6 sections with 0.5µm spacing were acquired and the presented images are a maximum projection of images deconvolved using softWoRx v4.1.2 inbuilt module. All conclusions based on imaging data were derived from at least three independent observations.

**Image processing and fluorescence intensity based quantifications** were performed in ImageJ (NIH, Bethesda, USA). Supplemental movies were converted from TIFF to MOV format using the inbuilt MPEG-4 compression. For overnight time-lapses that exhibited drifting, presented images were aligned using the MultiStackReg1.45 Plugin. All quantifications were performed on raw images.

**The lifetime of the fusion focus** was measured on time-lapse images as the interval between the fluorescently labeled marker proteins first focalizing^16^ and cell fusion. Results are reported with the box-and-whiskers plot where center lines show the medians, box limits indicate the 25th and 75th percentiles, whiskers extend 1.5 times the interquartile range from the 25th and 75th percentiles, outliers are represented by dots and the Kruskal-Wallis test p-value is reported.

To distinguish the expression of *mei3* from P- and M-genome we performed crosses between heterothallic strains where only one partner carried the *mei3-sfGFP* allele, while the other partner had the unlabeled, wildtype gene. **Quantification of Mei3-sfGFP fluorescence** induction used cell fusion as time zero, evidenced as the first timepoint with exchange of cytosolic RFP between partners. The mean signal in individual partners prior to fusion, and thus prior to *mei3* induction, was considered as background signal. Mean fluorescence of the whole zygote was recorded over time and the average of the indicated number of cells is reported with shaded regions denoting mean ± standard deviation. The post-fusion time for the mean zygotic intensity to reach 50 arbitrary units above background was recorded for each zygote and the results are reported with the box-and-whiskers plot where center lines show the medians, box limits indicate the 25th and 75th percentiles, whiskers extend 1.5 times the interquartile range from the 25th and 75th percentiles, outliers are represented by dots and the Kruskal-Wallis test p-value is reported.

For **Mi-sfGFP nuclear entry dynamics** we used the last time-point prior to exchange of Mi-sfGFP between partners as time zero. The nuclei of the zygote were clearly identifiable based on Mi-sfGFP signal alone, which allowed us to outline the nuclear region required in these quantifications. The signal in the P-cell prior to cell fusion was considered as background. Mean fluorescence of each nuclear region was recorded over time and the average of the indicated number of cells is reported with shaded regions denoting mean ± standard deviation and the Kruskal-Wallis test p-value is reported.

For **quantification of dynamics of NLS-sfGFP-SynZip4, sfGFP-SynZip4 and mCherry-SynZip3** we used the last time-point prior to exchange of red fluorophore between partners as time zero. The nuclear region was outlined based on the accumulation of the fluorescent signals in the central cell region. The signal in the partner cell lacking the fluorophore prior to cell fusion was considered as background signal. Mean fluorescence of each nuclear region was recorded over time and the average of the indicated number of cells is reported with shaded regions denoting mean ± standard deviation and the Kruskal-Wallis test p-value is reported. Note that the mCherry-SynZip3-expressing cell also displays background vacuolar fluorescence, likely due to pre-fusion degradation of some mCherry-SynZip3.

---

**Figure S1.**
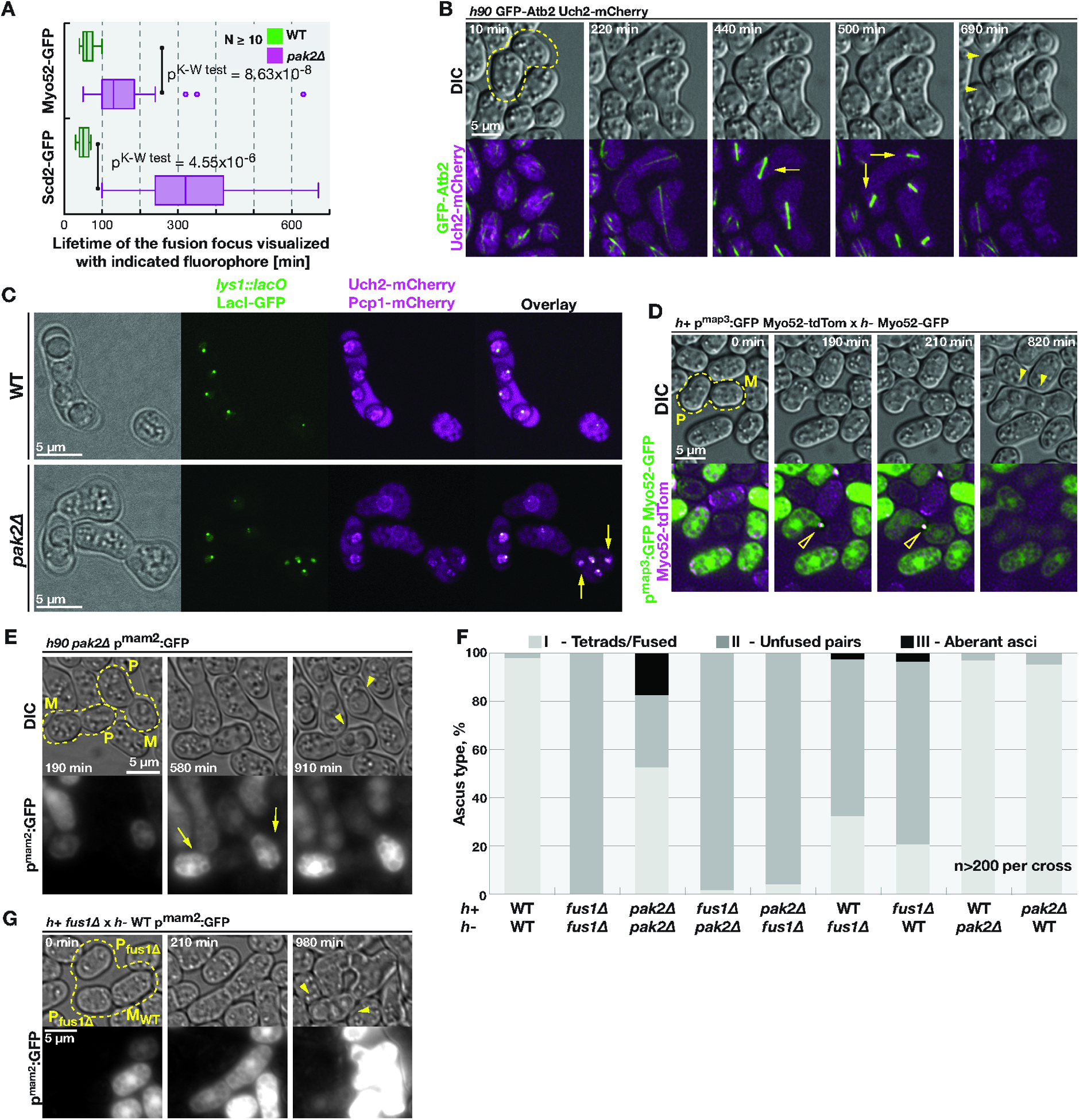
Transient cell fusion results in ectopic meiosis in the P-cell. **(A)** Box-and-whiskers plot (see Methods) reports the lifetime of the fusion focus visualized by the indicated fluorophore in homothallic wildtype and *pak2Δ* mating cells. Kruskal-Wallis test p^value^ is reported**(B)** Wildtype control for data presented in Fig. 1B. Time-lapse of homothallic wildtype mating cells expressing GFP-α-tubulin in green and nuclear marker Uch2-mCherry in magenta. Note karyogamy in the outlined mating pair, the meiotic spindles (arrows) and spores (arrowhead). **(C)** Micrographs of cells in mating mixtures of homothallic wildtype and *pak2*Δ strains with *lys1* chromosomal loci labeled by lacO:GFP-NLS-LacI system in green and nuclei visualized by Uch2-mCherry in magenta. Arrows point to spores lacking the *lys1* locus. **(D)** Wildtype control for data presented in Fig. 1D. Time-lapse of homothallic pak2Δ mating cells expressing cytosolic GFP from the P-cell specific *p*^*map3*^ promoter. Note formation of the fusion focus (empty arrowhead), labeled by Myo52-GFP in the M-cell and Myo52-tdTomato in the P-cell, followed by exchange of cytosolic GFP and spore formation (full arrowheads). **(E)** Time-lapse showing mating of homothallic *pak2Δ* cells expressing GFP from the M-cell specific *p*^*mam2*^ promoter. Note that the GFP exchange is followed by fusion pore closure and build up of the GFP signal only in the M-cells (arrows) while P-cells proceed to form spores (arrowheads). **(F)** Heterothallic strains of indicated genotypes were mixed and induced to mate on media plates. Two days later we quantified frequencies of phenotypes indicated in the insets of Figure 1A. **(G)** Time-lapse showing mating of *h+ fus1*Δ mutant and *h-* wildtype cells expressing GFP from the M-cell specific *p*^*mam2*^ promoter. Note that transient cell fusion between indicated partners is followed by resealing of the fusion pore, as visualized by continued accumulation of GFP only in the M-cell, and sporulation (arrowheads) in the P-cell.

**Figure S2.**
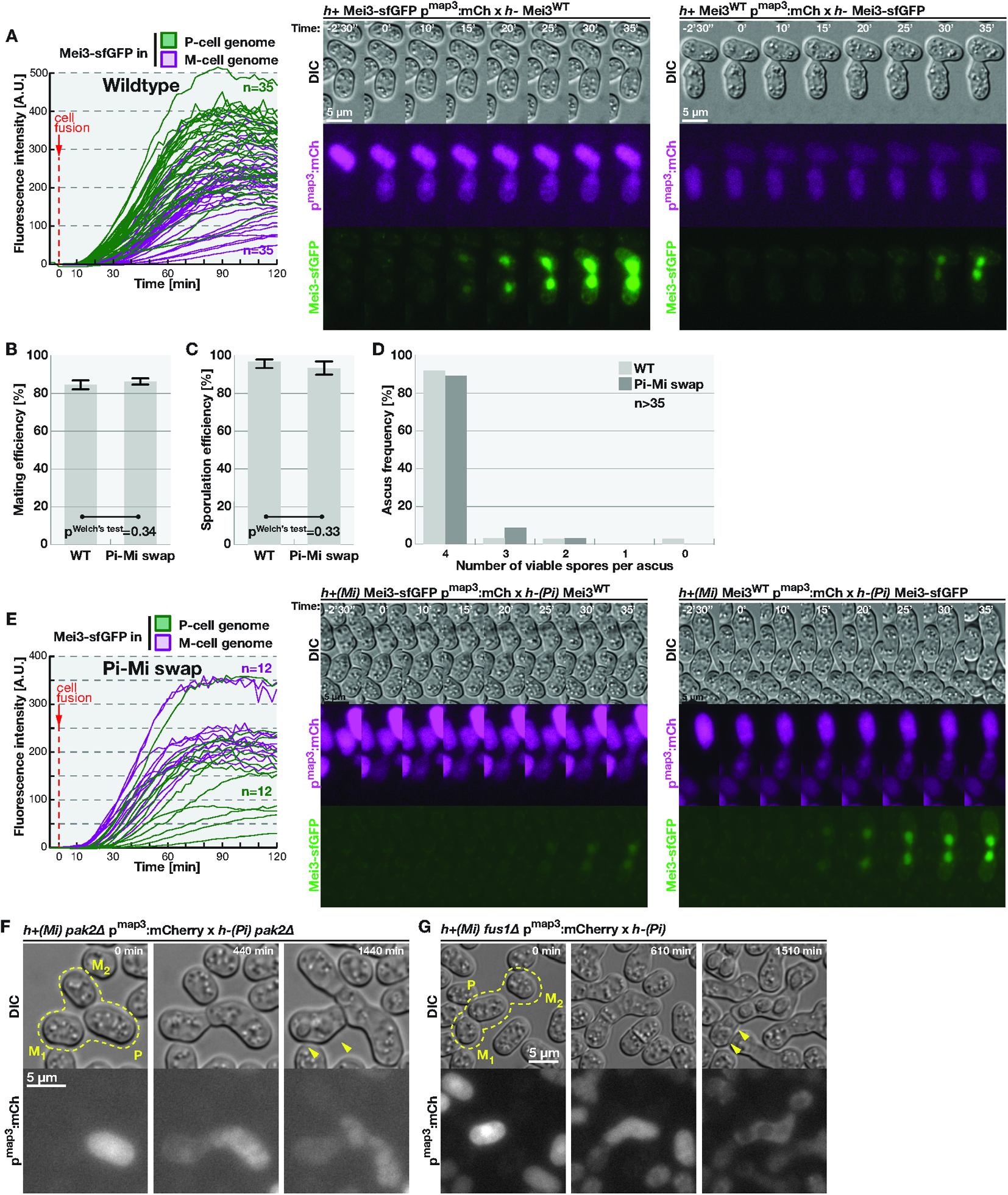
Induction of the meiotic inducer Mei3 occurs first from the genome of the cell expressing homeodomain protein Pi. **(A)** Left panel shows individual traces quantifying fluorescent signal from zygotes of heterothallic wildtype cells where only one partner has *mei3* tagged with sfGFP (see Figure 2C for average curves). Right panels show time-lapse of wildtype cells with P-cells expressing cytosolic mCherry and, as indicated, mei3 is tagged with sfGFP in only one partner. Note more rapid signal induction when Mei3 was labeled in the P-cell. **(B,C)** Quantification of mating and sporulation efficiencies in heterothallic wildtype and strains with swapped Pi and Mi coding sequences induced to mate on MSL-N agar plates for 28 hours. **(D)** Asci produced by mating heterothallic wild type or strains with swapped Pi and Mi were subjected to tetrad dissection and the number of spores that developed colonies counted. **(E)** Left panel shows individual traces quantifying fluorescent signal from zygotes of heterothallic cells with swapped Pi and Mi expression where only one partner has *mei3* tagged with sfGFP (see Figure 2D for average curves). Right panels show time-lapse of mating heterothallic strains with swapped Pi and Mi where *h+* cells express cytosolic mCherry and, as indicated, *mei3* is tagged with sfGFP in only one partner. Note more rapid signal induction when Mei3 was labeled in the M-cell. **(F)** Heterothallic *pak2Δ* strains with swapped Pi and Mi coding sequences and expressing P-cell specific p^map3^:mCherry were induced to mate. Transient fusion, visualized by florescence exchange, leads to haploid meiosis in the M1-partner (arrowheads indicate spores). Note that P-cell proceeds to attempt to mate with M2-cell. **(G)** Time-lapse showing mating cells with swapped Pi and Mi coding sequences. The P-cell also lacks *fus1* and expresses cytosolic mCherry. Note the spores forming in the M1-cell upon transient fusion (arrowheads) and that the P-cell subsequently mates with M2-cell.

**Figure S3.**
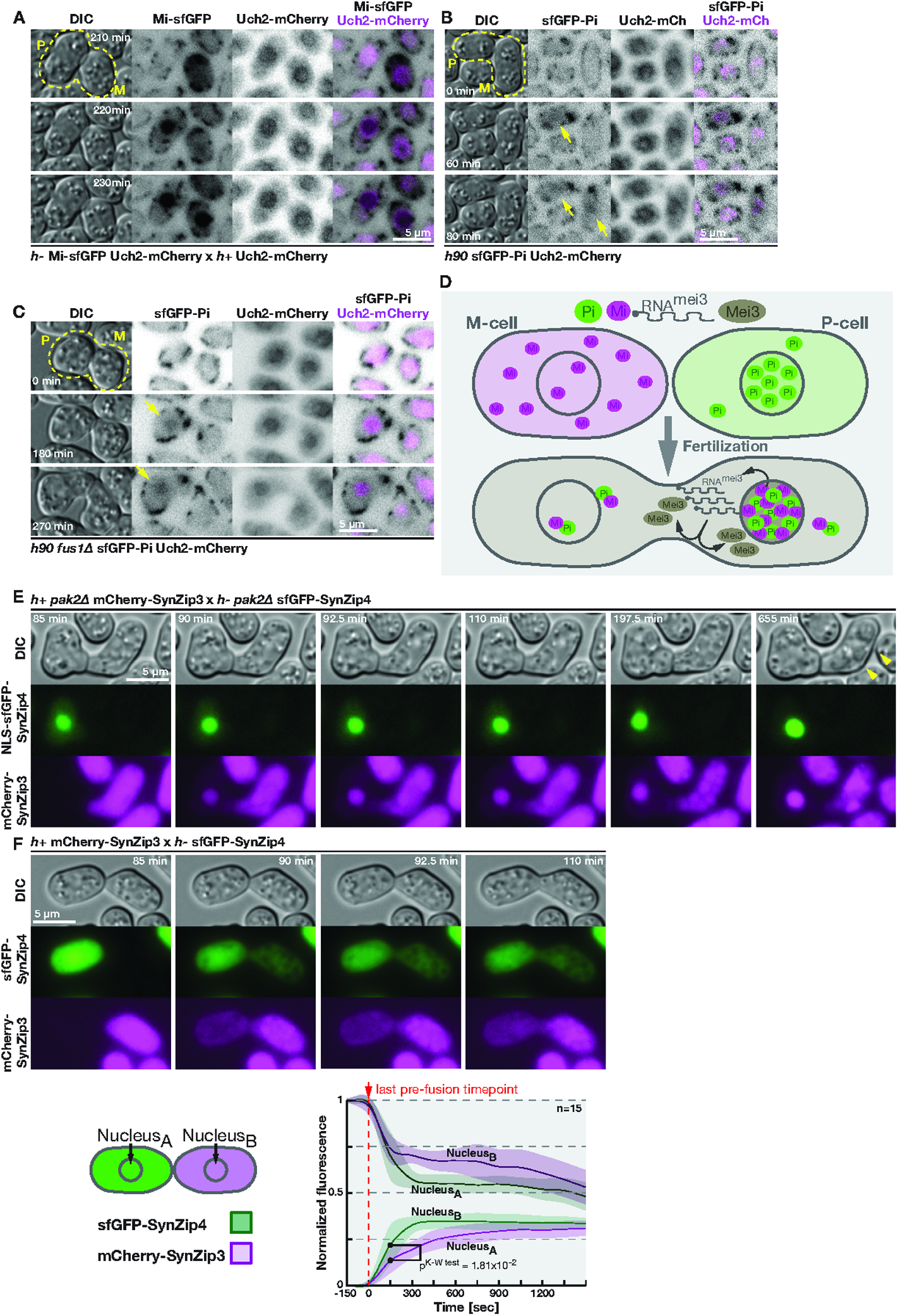
Meiotic regulators Pi and Mi exhibit asymmetry in localization in early zygotes. **(A)** Time-lapse showing mating of heterothallic cells expressing nuclear marker Uch2-mCherry (magenta) with the M-cell co-expressing sfGFP tagged endogenous Mi. Note the cytosolic Mi-sfGFP signal in M-cells that rapidly accumulates in the P-cell nucleus after partner fusion. Note that the dotty signal at cell cortex in the green channel is background fluorescence, likely produced by mitochondria. **(B)** N-terminally sfGFP-tagged endogenous Pi exhibits a weak nuclear staining in P-cells during mating in homothallic cells co-expressing nuclear marker Uch2-mCherry (magenta). Arrows point to nuclear sfGFP-Pi signal in the P-cell (middle panel) and the zygote (bottom panel). **(C)** Time-lapse showing homothallic, fusion defective, *fus1*Δ cells co-expressing N-terminally sfGFP-tagged endogenous Pi and nuclear marker Uch2-mCherry. Arrows point to Pi nuclear accumulation. **(D)** Model for the asymmetric localization of the bi-partite transcriptional activator produced by Pi and Mi in early zygotes, which results in asymmetric expression of *mei3* between parental genomes. **(E)** Time-lapse showing transient cell fusion between *pak2Δ* cells expressing NLS-sfGFP-SynZip4 and mCherry-SynZip3. Note accumulation of both fluorophores in the M-nucleus and minimal delivery of the green fluorophore into the P-cell that eventually sporulates (arrows) **(F)** Top panel follows the mating of wildtype cells expressing cytosolic sfGFP-SynZip4 and mCherry-SynZip3. Bottom panel quantifies both fluorescent signals in the central region of the two cells corresponding to the two nuclei, as labeled on the scheme, and is presented as in Fig. 3F.

**Figure S4.**
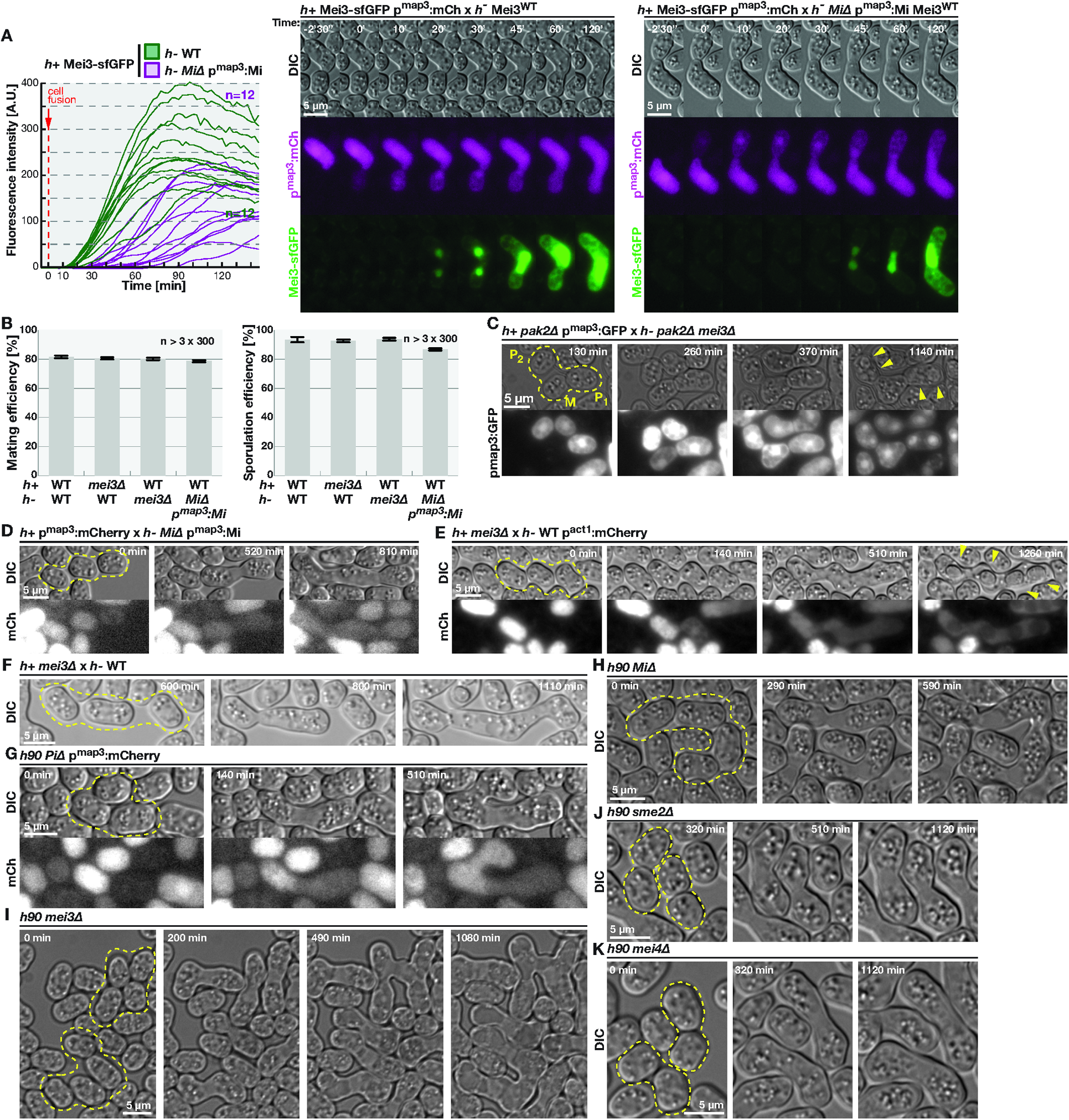
Rapid induction of Mei3-Mei2 signaling is required to suppress mating in zygotes and prevent polyploid formation. **(A)** Left panel shows the quantification of the green fluorescent signal in zygotes produced by mating either wildtype or Δ*Mi p*^*map3*^:*Mi* M-cells with P-cells encoding Mei3-sfGFP. Profiles of individual cells are shown (see Figure 4A for average curves). Right panels show time-lapse of P-cells expressing cytosolic mCherry (magenta) and Mei3-sfGFP (green) mating with M-cells that are either wildtype or express Mi from the P-cell specific p^map3^ promoter. Note delayed Mei3-sfGFP signal when Mi expression is delayed**. (B)** Heterothallic strains with indicated genotypes were mixed and induced to mate on MSL-N agar plates. Charts report mating and sporulation efficiencies quantified after two days incubation, p^Welch’s^ ^test^ > 0.05 for all comparisons to wildtype. **(C)** Time-lapse shows *pak2Δ* P-cells expressing cytosolic GFP mating with *pak2Δ mei3Δ* double mutant M-cells. Note that transient cell fusion is followed by spore formation (arrowheads) in both indicated P-cells. **(D)** Time-lapse showing that mating of M-cells with Mi expressed solely from the p^map3^ promoter with wildtype P-cells expressing cytosolic mCherry results in zygotes undergoing fusion with additional partners evident as exchange of the cytosolic fluorophore. **(E-F)** Time-lapse showing that zygotes undergo repeated cell fusion, evident from exchange of cytosolic mCherry, in crosses where *mei3* was specifically deleted in the P-cells. **(G-I)** Time-lapse showing re-fusion of PiΔ, MiΔ and mei3Δ homothallic cells, as indicated. **(J-K)** Time-lapse showing diploid zygote formation in sme2Δ and mei4Δ homothallic cells, as indicated. Note that zygotes do not exhibit any growth or attempt mating with other cells.

**Figure S5.**
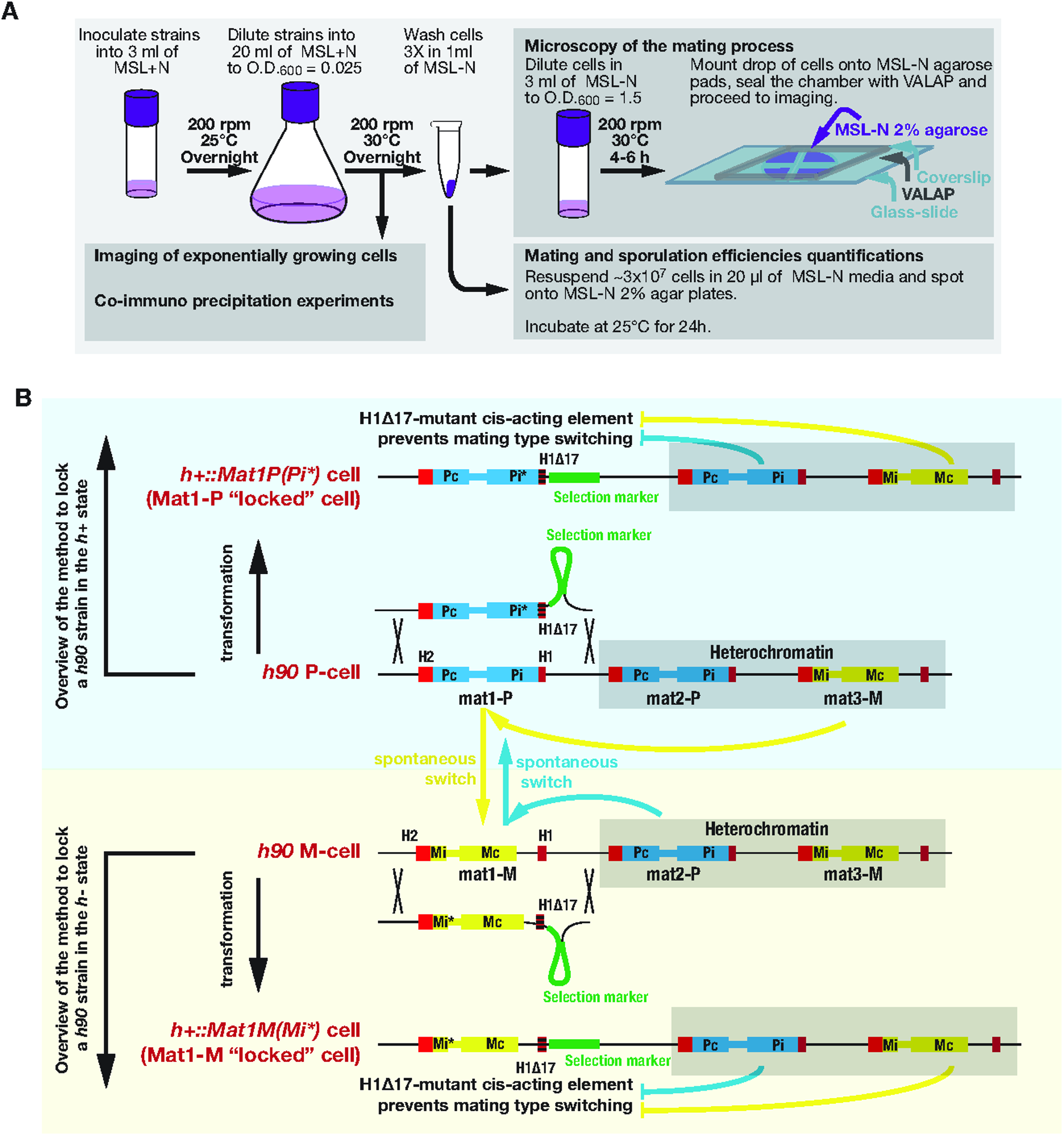
Schematic overview of growth conditions and *mat1* locus manipulations. **(A)** Overview of growth conditions used in this study. See Methods section for details. **(B)** Schematic representation of *mat1* locus manipulations. The wildtype homothallic P and M-cell *mat* loci are presented in the middle of the scheme. Red boxes denote H2 and H1 homology boxes identical between the three *mat* loci. Blue boxes represent Pc and Pi genes expressed exclusively in the P-cell and encoded by the *mat2* locus while yellow boxes denote Mc and Mi genes encoded by the *mat3* locus and expressed in M-cells only. Gene expression from *mat2* and *mat3* loci is inhibited by heterochromatin state (gray box). The sequences at the *mat1* locus are derived from sequences at the other two *mat* loci during spontaneous mating type switching (blue and yellow arrows) which relies on the H1 and H2 homology boxes. Transformation with a DNA fragment carrying the mutant H1Δ17 homology box (shaded red box) results in cells unable to switch their mating type.

**Figure S6.**
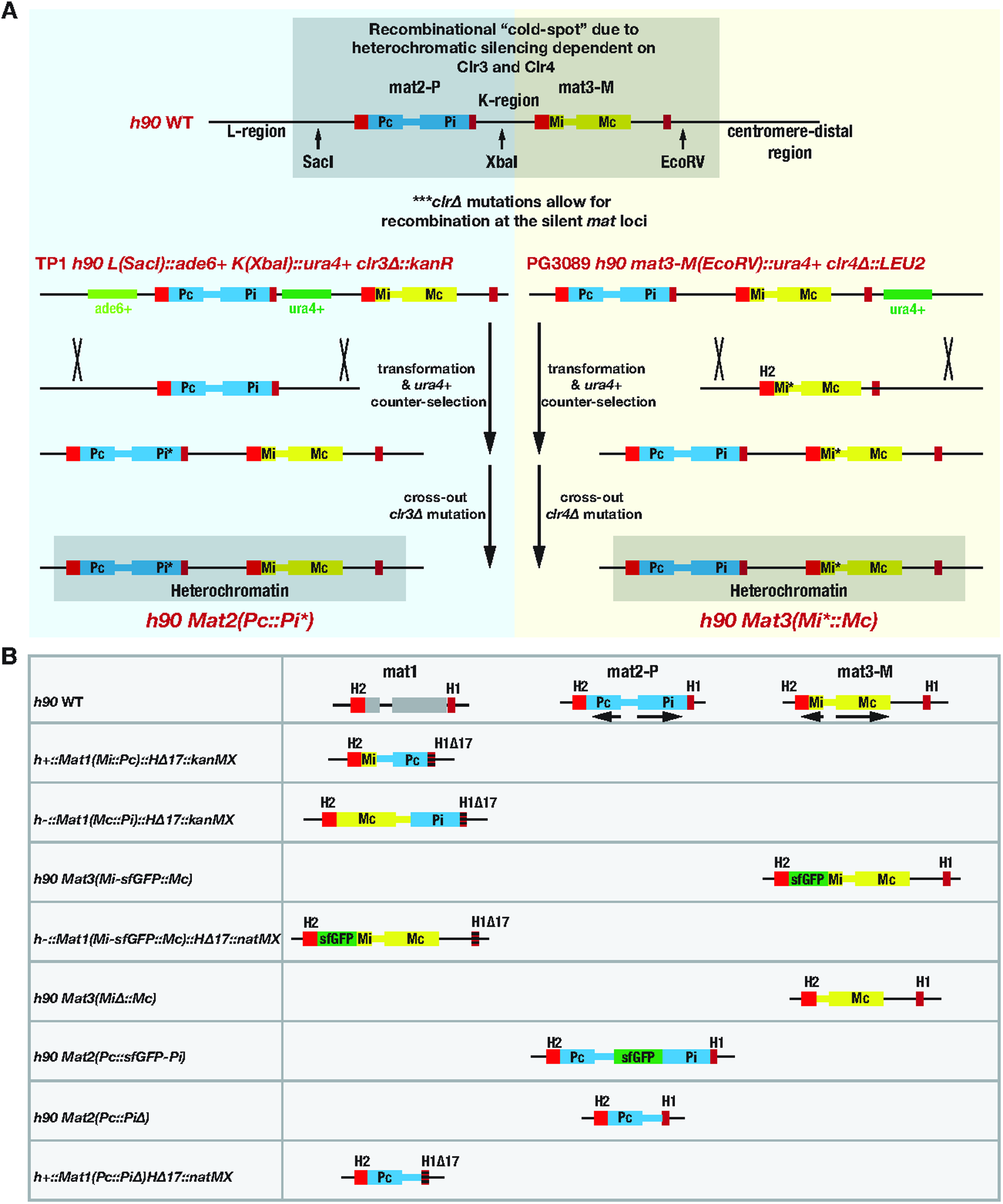
Schematic overview of *mat2* and *mat3* loci manipulations and obtained mat loci mutants. **(A)** Schematic representation of *mat2* and *mat3* loci manipulations. Wildtype homothallic strain is presented on the top. Red boxes denote H2 and H1 homology boxes identical between the three *mat* loci. Blue and yellow boxes represent genes encoded by the *mat2* and *mat3* loci respectively. Recombination at the *mat2* and *mat3* loci is inhibited by heterochromatin state (gray box). Restriction enzyme sites denote positions where prototrophic genes have been integrated in the TP1 and PG3089 strains depicted bellow which are used to manipulate the *mat2* and *mat3* loci respectively. Ablation of genes necessary for heterochromatin formation allows targeting the *mat2* and *mat3* loci by homologous recombination. **(B)** Schematic representation of *mat* loci mutants obtained in this study. Color-coding is as in Fig. S5B.

**Table S1:**
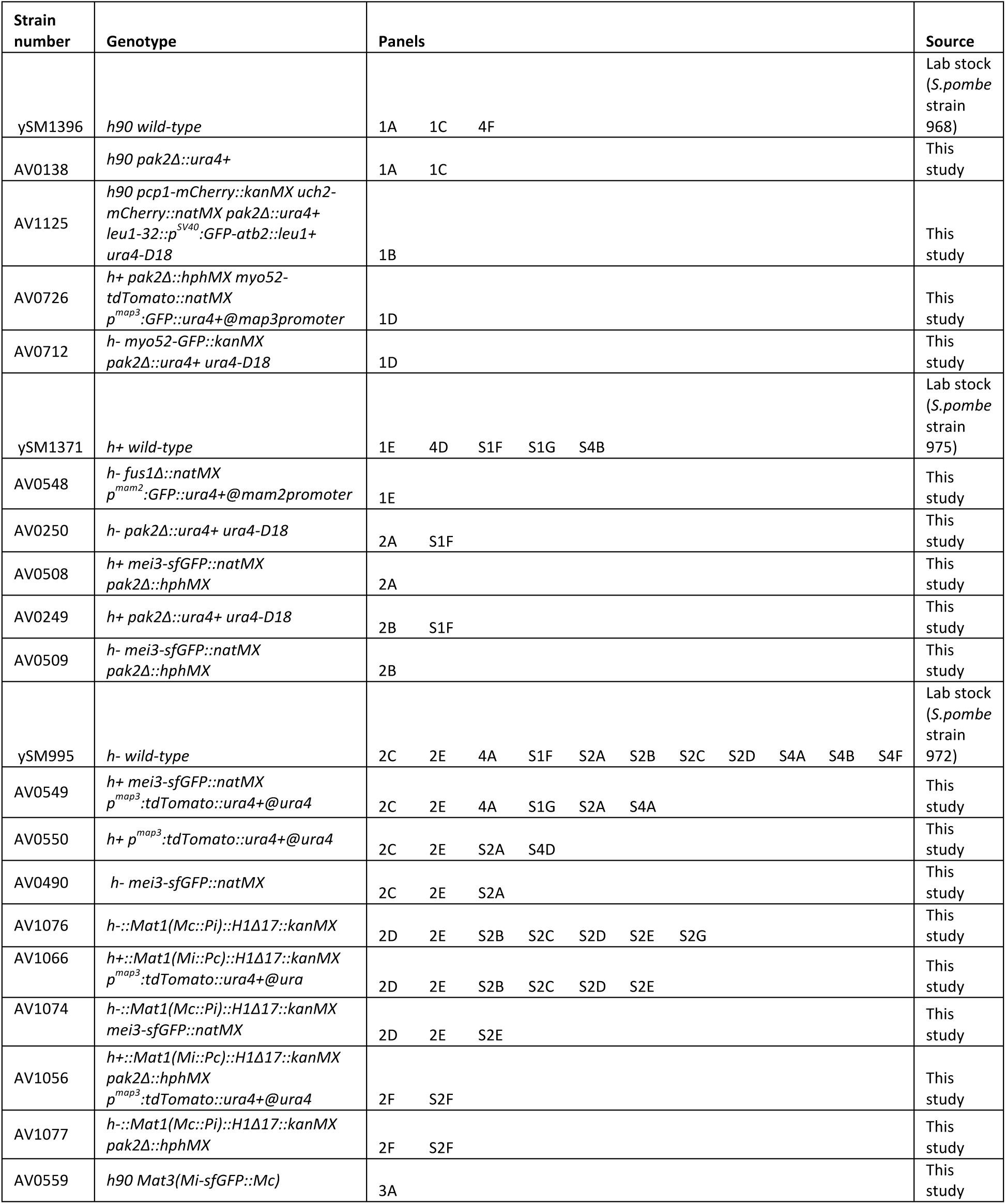
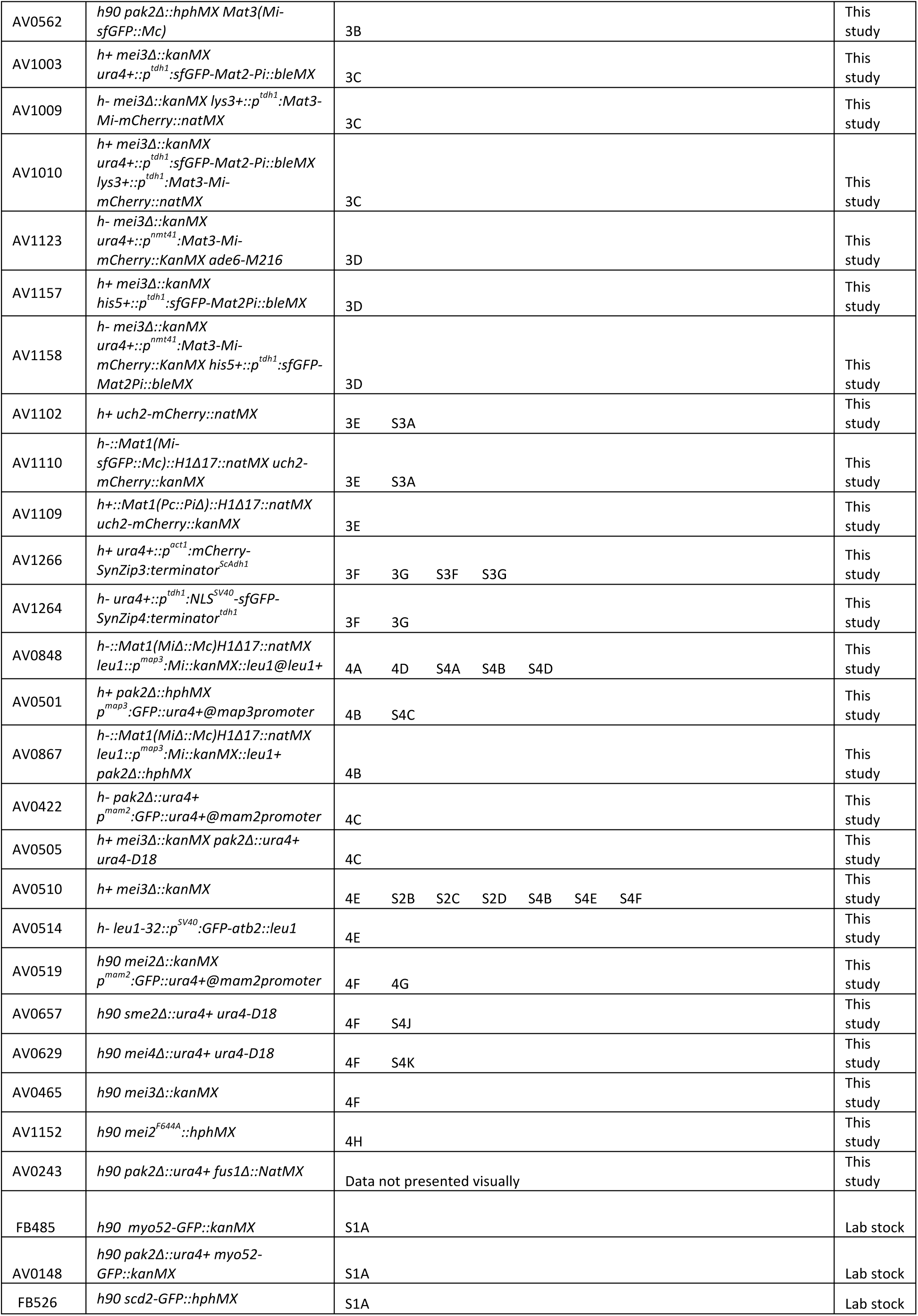
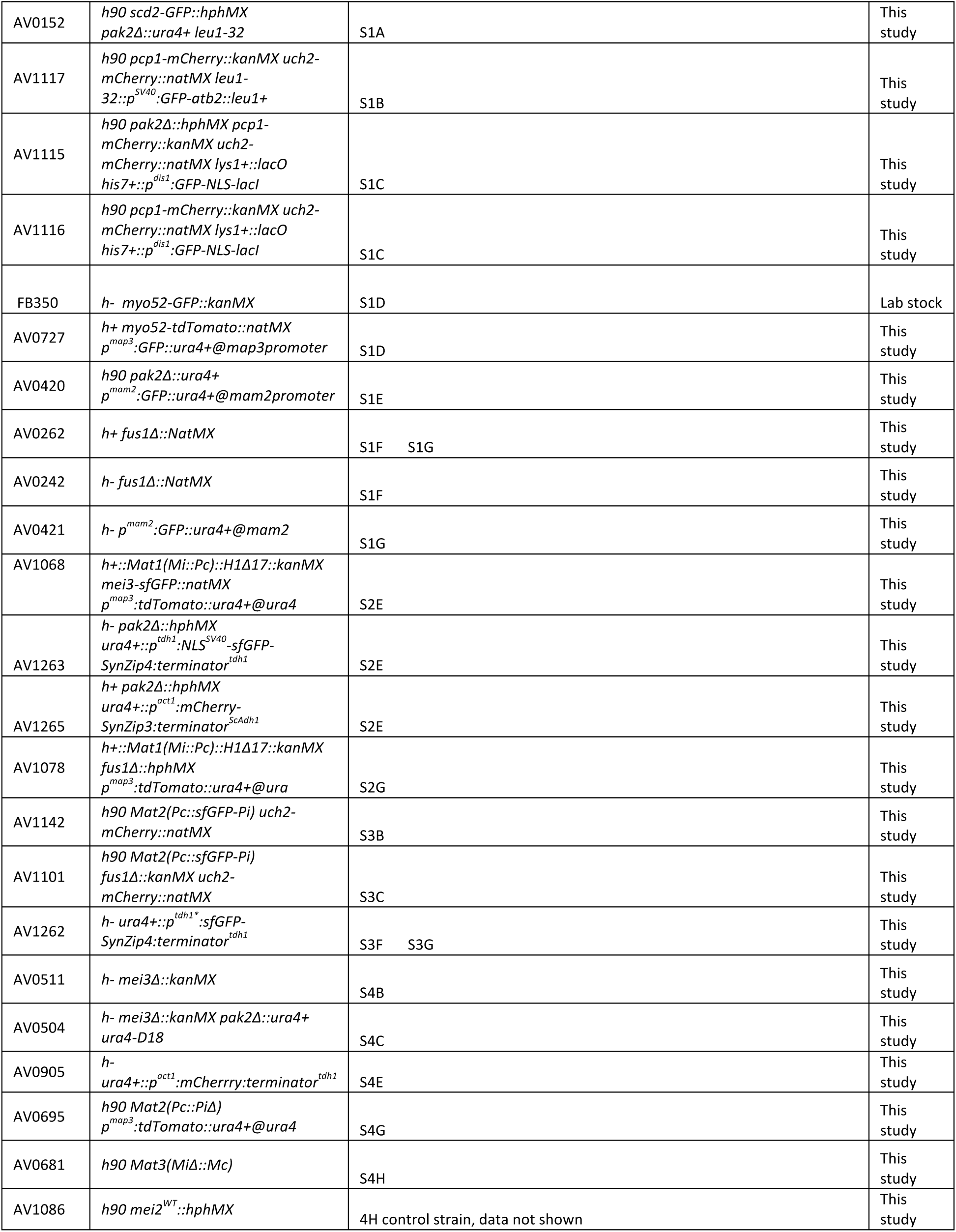
Strain list

**Table S2:**
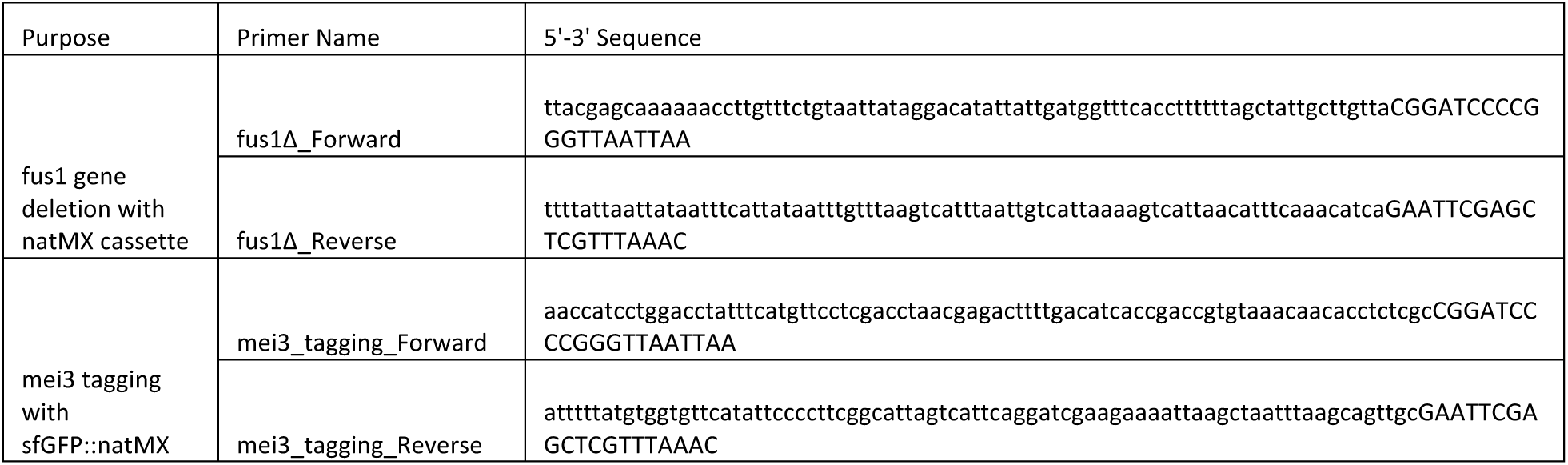
Primer list

